# An individual-based simulation framework exploring the ecology and mechanistic underpinnings of larval crowding in laboratory populations of *Drosophila*

**DOI:** 10.1101/2023.07.30.551144

**Authors:** Srikant Venkitachalam, Amitabh Joshi

## Abstract

The study of larval competition in laboratory populations of *Drosophila*, implemented via the crowding of larval cultures, has contributed greatly to the understanding of the ecology of competition, the evolution of larval competitive ability, and formed the basis of rigorous testing of the theory of density-dependent selection. Earlier studies led to the view that the outcomes of larval competition, and resulting evolutionary consequences of crowding-adaptation, could largely be understood by varying the starting density of individuals in a crowded culture. However, recent studies have shown that the results of adaptation to larval crowding may not be well predicted by the total larval density (i.e., total starting individuals/total volume of food). Cultures raised at the same total density but at different egg number and food volume combinations were shown to have different underlying density-specific fitness functions, and crowding-adaptation in each of these cultures was attained through different evolutionary trajectories as well. A recent study showed that cultures with not just the same density, but the same egg and food volume combination, achieved through food columns of differing diameter and height, could also differ greatly in fitness-related trait outcomes. In that study, the density of larvae in the feeding band (volume of food close to the surface in contact with air, to which larval feeding is largely restricted) was a very important factor in predicting the outcomes of larval competition. Given these recent findings, it is important to understand the overall role of feeding band density, and how it influences density-specific fitness functions in different kinds of crowded cultures. As the older models of larval competition are now insufficient to capture current empirical data, we constructed an individual-based simulation framework informed in part by these more recent findings, in order to better understand the evolutionary ecology and mechanistic underpinnings of larval competition, and predict robust experiments for expanding our understanding of the process of larval competition in *Drosophila*.

## Introduction

The theory of density-dependent selection, first formally described by MacArthur (1962) and MacArthur and Wilson (1967), is a vital interface between the evolutionary and ecological approaches to competition (reviewed in Mueller, 1997, 2009; Joshi et al., 2001). At its core, it posits that fitness landscapes associated with various traits can change across population densities – i.e., the relationship of a trait variant with Darwinian fitness could depend on the population density it experiences.

Over the last few decades, many theoretical and experimental studies have aimed to examine the evolution of traits under different population density conditions (reviewed in Mueller, 1997; Joshi et al., 2001). On the experimental side, some of the most rigorous and detailed work has been carried out through selection experiments using outbred laboratory *Drosophila* populations (reviewed in Sarangi et al., 2016; Venkitachalam et al., 2022). Adaptation to chronically crowded conditions has been used to explore the evolution of traits under high-density selection compared to respective low-density reared control populations.

The earliest studies focused on crowding populations at all life-stages through serial transfers (Mueller and Ayala, 1981). While this approach yielded some insights into the evolution of traits under high density conditions, crowding experienced at both larval and adult stages was a confounding factor when drawing inferences (Mueller et al., 1993). Thus, later studies focused largely on adaptation to larval crowding (but see also studies on adaptation to adult crowding by Joshi and Mueller 1997; Joshi et al., 1998). These larval crowding studies have previously been described in detail (reviewed in Prasad and Joshi, 2003; Sarangi et al., 2016; Venkitachalam et al., 2022). The overall results from these studies were the following –

- Larval competitive ability evolves to be greater in crowding-adapted populations as compared to the low-density reared controls, under most selection studies carried out so far (Mueller, 1988a; Nagarajan et al., 2016; Sarangi et al., 2016; Sarangi, 2018; Venkitachalam et al., 2023a; see Joshi and Mueller (1996) for an indirect inference of the same).
- The exact details of initial egg number, food volume and container dimensions (usually a cylindrical glass vial) determine which competitive ability linked traits evolve, and to what extent, in a given crowded scenario (Sarangi, 2018; Venkitachalam et al., 2022, 2023a).
- Even under the same high-density conditions implemented via different combinations of initial egg numbers and food volume, the outcome of crowding with respect to fitness-related traits can be different, and this can also lead to the evolution of different traits under each condition (Sarangi, 2018).

Thus, it became apparent that a better understanding of the ecology of larval competition under different kinds of crowded scenarios would be vital to better understanding the evolution of larval competitive ability.

In parallel with, and also preceding the studies on evolution of adaptations to crowding, there was a long tradition of experimental and theoretical studies concerned with the ecology of larval competition in *Drosophila*. These studies ranged from describing the general outcome of larval crowding in *Drosophila* (Sang, 1949; Chiang and Hodson, 1950; Bakker, 1961; Ohnishi, 1976), to the role of waste in crowded conditions (Weisbrot, 1966; Botella et al., 1985), the cessation of developmental activity (Ménsua and Moya, 1983; González-Candelas et al., 1990),and the measurement of larval competitive ability in competition experiments between various strains (Bakker, 1961, 1969; Gale, 1964; Kearsey, 1965; Seaton and Antonovics, 1967; Mather and Caligari, 1981; de Miranda et al., 1991; Santos et al., 1992). Among these, studies by K. Bakker were particularly detailed, and inspired several models to understand and predict the results of larval competition, in addition to a verbal model described by Bakker himself (Bakker, 1961). These models, which at least in part covered the outcomes of larval competition, ranged from verbal (Bentvelzen, 1964) to mathematical (De Jong, 1976; Nunney, 1983; Mueller, 1988b; Jansen and Sevenster, 1997; Tung et al., 2019). In each of these models, which were based directly on Bakker’s data or from results of similar experiments, the primary end point of a culture experiencing competition under extreme crowding was the running out of food – and the primary outcomes were the differences in survivorship and body size of various types. These models largely considered different numbers of ‘larvae’ competing for some limited quantity of food, thereby varying larval density. Moreover, most of these models were concerned with describing or predicting the long-term consequences of the overall results of larval crowding as a whole. The understanding that the details of crowding experienced at similar overall densities, but at differing combinations of egg number and food volume, could result in different outcomes of fitness-related traits was largely absent. Also missing from these earlier models were the effects of metabolic waste products, which could impact the outcomes of competition along the axis of pre-adult development time in addition to survivorship (but see an independently-developed model by Moya and Castro (1986) for some consideration of the effects of metabolic waste on larval competition).

Adding to these unconsidered details, a recent study conducted by us decomposed the effects of larval crowding along the axes of starting egg number, height of the food column and the cross-section surface area of the cylindrical container (Venkitachalam et al., 2023b). This was largely inspired by the studies conducted by J.H. Sang (Sang, 1949). We found that the same combination of egg number and food volume cast into cylinders of different dimensions, via different food column height and surface area combinations, could greatly affect the egg to adult survivorship and the distribution of development time in a culture. Furthermore, we found that the density of larvae at the ‘feeding band’ – a shallow volume of food in contact with air, to which larval feeding is restricted – predicts the outcome of larval crowding better than the total density, as described by the starting number of eggs divided by the total starting volume of food.

In light of the recent advances in our understanding of the nuances of larval competition and of the adaptation to chronic larval crowding, there is a need to mechanistically explore how the differences in food surface area and food column height may be leading to differences in vital fitness-related traits such as survivorship and development time. Exploring this whole range of possible combinatorial scenarios experimentally is a daunting task from a logistical perspective. Therefore, we have developed an individual-based simulation framework that explores the mechanisms by which the outcomes of larval crowding are ultimately mediated. Such a model that predicts the results of larval competition under realistic scenarios can also be used to explore the consequences of long-term selection for adaptation to larval crowding under different conditions, further refining our understanding of density-dependent selection. This aspect has been elaborated in the discussion section (section 4.4).

The simulation framework described in the current study attempts to capture the processes of larval growth and the varied ways in which competition may actually occur in a crowded culture. This work is a follow up and expansion of a simple novel model developed by us and published as a Master’s thesis by one of the authors (Venkitachalam, 2017). The current study limits the simulation to a single generation of a *Drosophila* larval culture, studying the outcome in terms of the distributions of fitness-related traits from various kinds of crowded cultures drawn from our recent empirical study (Venkitachalam et al., 2023b). The simulation framework is useful inasmuch as it may be helpful in generating predictions, confronting hitherto unconsidered mechanisms and directing precise experiments in situations wherein general exploratory experiments may be logistically daunting.

### 1. Traits involved in larval competition

Before setting up an individual-based simulation on larval competition, we list the traits being imparted to every competing individual. Each of these traits has been, or can potentially be, implicated in some way to be important to larval competitive ability – either in empirical studies involving the evolution of increased larval competitive ability, or in some form of logical argument. For this listing of traits, we describe a heuristic model for understanding the importance of various traits to larval competition, across three distinct time-periods of a culture.

#### 1.1. A heuristic model of larval competition in a crowded culture

We imagine a crowded culture wherein we single out one individual (fig. 1.a). Over the timespan of the culture, this individual starts in the culture as an egg, which hatches into a larva. This larva then feeds on the available food and grows (through three instars), in competition with the myriad other larvae competing for the same limited resources. Ultimately, the crowded culture starts running out of food and/or building up toxic concentrations of metabolic wastes. In order to survive, the focal larva must achieve at least its minimum critical mass for pupation by feeding on a minimum amount of food (Beadle et al., 1938; Chiang and Hodson, 1950; Bakker, 1959, 1961; Robertson, 1963) before the food runs out and/or the ambient waste concentration becomes lethal for further food consumption.

**Figure 1.**
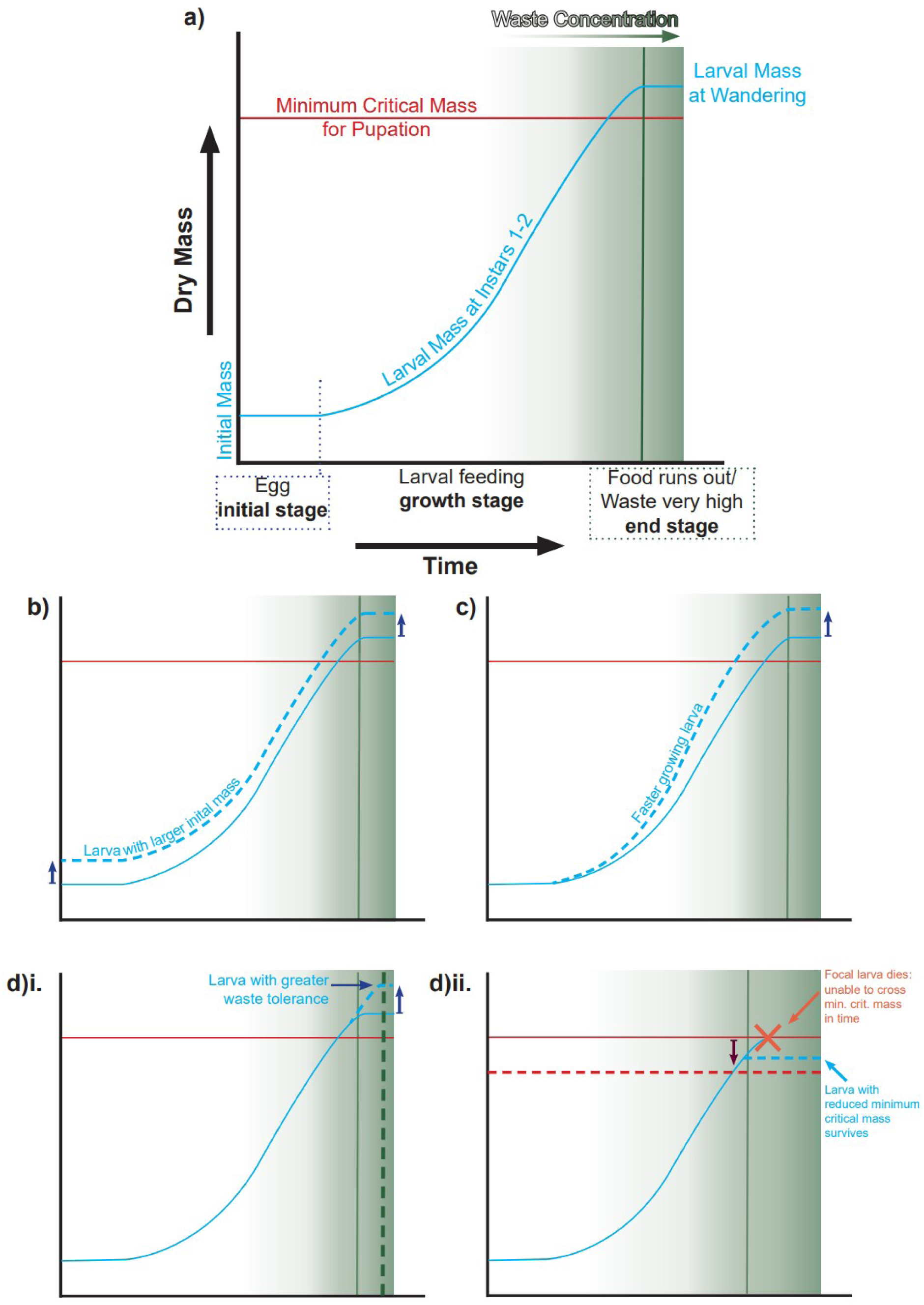
Heuristic model of larval competition (see introduction section 1.1 for more details). a) This plots the growth profile of a single imagined individual from an imagined crowded *Drosophila* larval culture. The individual can go through three fairly distinct stages through the life of the crowded culture. It begins in the initial stage as an egg. The time of hatching determines the beginning of the growth stage of the larva. As multiple larvae feed in a crowded culture, food starts running out and/or metabolic waste concentration starts building up, as highlighted by a green background. This determines the end stage of the culture for the larva, when it must leave the system due to food running out or waste toxicity becoming too high, denoted by the solid green line. In order to commit to wandering stage and become an adult, the larva must cross its minimum critical mass required for pupation, else it dies. This threshold mass is drawn as a red line. b) Traits that may improve competitive ability at the initial stage (see Table 1). Besides the focal individual (solid blue line), we imagine another competing individual (dashed blue line). This second individual has a larger initial larval mass, achieved via greater egg mass, compared to the focal individual. All else being equal, the individual with the larger egg mass will have a greater body mass than the focal larva by the end stage, giving it a potential competitive edge. An imagined individual with a faster egg hatching time would get a similar competitive edge, by feeding earlier and effectively having a larger mass by the time the focal individual hatches. c) Traits that may improve competitive ability at the growth stage (see Table 2). We imagine another individual (dashed blue line) with a greater growth rate of larval mass than the focal larva (solid blue line). All else being equal, the individual with the greater growth rate will have a greater body mass than the focal larva by the end stage, giving it a potential competitive edge. d) Traits that may improve competitive ability at the end stage (see Table 3). At this stage, a second larva (dashed blue line) can be more competitive than the focal larva (solid blue line), all else being equal, via two distinct ways:

i. Increased waste tolerance – the larva with the higher waste tolerance can continue feeding in greater concentrations of waste (dashed green line denotes the increased waste tolerance threshold), eclosing later but at a greater mass.
ii. Decreased minimum critical mass to pupation – the larva with lower minimum critical mass requirement, denoted by the dashed red line, can escape very high waste levels and become an adult, while a larva with a greater threshold requirement will succumb.

**Table 1.** Traits that may be important to competitive ability at the initial stage.

| Trait | Significance | Notes | Mean | c.o.v. | Source(s) |
| --- | --- | --- | --- | --- | --- |
| **i. Egg hatching time** | The egg hatching time determined the time step at which an egg hatched into a larva. | Each individual's hatching time was implemented as a countdown from the start of the culture time. | 20 hours | 1% | Venkitachalam et al., 2022 |
| **ii. Egg length** | This translated into the initial larval length. | Egg shell dimensions were not considered. | 0.5 mm | 5% | Venkitachalam et al., 2022 |
| **iii. Egg width** | This translated into the initial larval cross-section diameter. | Egg shell dimensions were not considered; diameter was used to calculate initial cross-section surface area of larva. | 0.18 mm | 5% | Venkitachalam et al., 2022 |

**Table 2.** Traits that may be important to competitive ability at the growth stage.

| **Trait** | **Significance** | **Notes** | **Mean** | **c.o.v.** | **Source(s)** |
| --- | --- | --- | --- | --- | --- |
| **i. Initial Bite Rate** | For a given bite mass, the initial bite rate determined the rate of food intake. | The bite rate increased in value across the larval growth period (see text). | 3000 bites per hour | 5% | Joshi and Mueller, 1996; Santos et al., 1997; Joshi et al., 2003 (c.o.v.); Sarangi et al., 2016 |
| **ii. initial bite mass** | For a given bite rate, the initial bite mass determined the rate of food intake. | The bite mass decreased in value across larval growth period (see text). | 0.007 % of initial larval mass | None | Arbitrary |
| **iii. food to biomass conversion efficiency mean** | Determined the fraction of food eaten that was converted to growth in larval mass, each time step. | This determined the mean efficiency per larva across time steps. | 30% of food eaten in a given time step | 5% | (Only mean) Bakker, 1969; de Miranda and Eggleston, 1988 |
| **iv. Efficiency within larva c.o.v.** | Determined the variation across time steps in food to biomass efficiency for a larva. | The efficiency of a larva per time step was randomly drawn from a normal distribution with mean as iii, and c.o.v. determined by iv. | 5% | 5% variation in c.o.v. across larvae | Inspired by de Miranda and Eggleston, 1988 |

**Table 3.** Traits that may be important to competitive ability at the end stage.

| Trait | Significance | Notes | Mean | c.o.v. | Source(s) |
| --- | --- | --- | --- | --- | --- |
| **i. Minimum critical (dry) mass to pupation** | Determined the dry body mass up to which a larva must grow in order to pupate. | Also determined maximum mass possible for a larva as: $3 \times \text{min. crit. mass}$ (largely based on patterns seen in previous data). | 180 $\mu\text{g}$ | 5% | Bakker, 1961; Santos et al., 1997 |
| **ii. Waste tolerance factor** | Determined the threshold of total dry mass of metabolic waste that could be non-lethally ingested by a larva. | The waste tolerance threshold of a larva at a given time was calculated as: the waste tolerance factor × current larval mass. | 0.095 | 10% | Arbitrary |

From the above description, we can imagine three (more or less) distinct time stages in the culture in which different traits can affect the focal larva’s competitive ability (fig. 1.a).

##### 1.1.1. Time stage 1: Initial stage of a culture

Assuming simultaneous placement of all eggs into the culture (see Jansen and Sevenster (1997) for a model without this assumption), all else being equal, the focal individual can have greater competitive ability by having a faster egg hatching time than other competitors, or by having a larger mass at hatching (fig. 1.b, discussed in Bakker, 1969). Both of these aspects increase the effective head start that the larva can have in its growth period (reviewed in Venkitachalam et al., 2023c). While a faster hatching time obviously confers greater head start, a larger initial larval mass reduces the amount of growth required by the larva to achieve its minimum critical mass (all else being equal), assuming larger eggs hatch into larger individuals. The evolution of both these traits has indeed been observed in multiple population sets adapted to respectively different details of larval crowding – each crowding adapted population, regardless of the exact conditions of larval crowding imposed, evolved greater egg length and egg width, as compared to the low-density control populations (Venkitachalam et al., 2022). Out of the three population sets studied, one set also evolved significantly reduced egg hatching time compared to controls, while the other two sets showed non-significant reductions (Venkitachalam et al., 2022).

##### 1.1.2. Time stage 2: larval growth period

In case of larvae feeding at the initial time window of a crowded culture, faster growth rate (i.e., rate of weight gain) should be paramount for survival. All else being equal, the focal larva would be competitively superior to other larvae by growing faster than them (fig. 1.c, discussed in Bakker, 1961). There are two primary ways by which growth rate could be affected:

###### 1.1.2.1. Modification of overall feeding rate

An increased rate of food ingestion, all else being equal, could drive faster larval growth rate. This could happen via either increased bite rate, measured as the number of bites taken per time step or increased bite size, leading to more food ingested per bite (Bakker, 1961; Robertson, 1963).

Bite rate has usually been termed ‘(larval) feeding rate’ in empirical studies, and has a rich history of study across laboratories and populations. It is measured as the total number of sclerite retractions per minute while feeding (Sewell et al., 1974; Joshi and Mueller, 1988). Initial studies on adaptation to larval crowding showed a positive relationship of feeding rate with larval competitive ability (Joshi and Mueller, 1988, 1996). Moreover, a selection study on increased feeding rate also resulted in the evolution of increased larval competitive ability (Burnet et al., 1977). This led to the general assumption that the evolution of increased larval feeding rate would also be associated with increased larval competitive ability (Joshi and Mueller, 1996; Borash et al., 2000). However, in later selection studies, crowding-adapted populations of *D*. *ananassae* and *D. nasuta nasuta* evolved increased larval competitive ability without evolving increased feeding rate (Nagarajan et al., 2016). The underlying explanation for this contrasting set of results was revealed to be dependent on the exact conditions of larval crowding imposed. *D. melanogaster* populations crowded at the same conditions as the *D. ananassae* and *D. nasuta nasuta* populations (different from the earlier *D. melanogaster* studies) also evolved greater larval competitive ability without increased feeding rate (Sarangi et al., 2016). Further complicating the relationship of feeding rate with larval crowding was a recent study carried out in our research laboratory by M. Sarangi (2018). The results from that study showed that crowding-adapted populations of *D. melanogaster* which had unchanged feeding rate compared to controls when assayed singly or under low-density conditions, showed the highest feeding rate in their native crowded conditions in vials (Sarangi, 2018). Thus, the plasticity of larval feeding rate across the crowding gradient has emerged as another potential factor that contributes to larval competitive ability, although further experimentation is required to elucidate the details and extent by which such plasticity manifests itself under differently crowded conditions.

Little is known about the bite size of *Drosophila* larvae, and likely even less about the dynamics and variance of bite size across different stages of larval growth under crowded conditions. A study by Robertson (1963) showed that mouth part size increased along with larval size across the larval growth period. We can thus assume bite size to also increase with larval size.

###### 1.1.2.2. Modification of food to biomass conversion efficiency

All else being equal, the focal larva could also enhance its growth rate by being more efficient in food utilisation than its competitors (Bakker, 1961). This efficiency could manifest itself in two possible ways – fraction of food converted to biomass, and time taken to convert the same quantity of food to biomass.

The importance of increased resource utilisation efficiency under high population density conditions has been stressed at least since the time density-dependent selection was formalised (MacArthur and Wilson, 1967; but see Joshi et al. (2001) for a contrasting view). However, results from experiments on larval crowding adaptation have yielded mixed results. Earlier selection studies indicated that crowding-adapted populations had evolved reduced efficiency with respect to minimum food required for pupation, indicating a reduction in the fraction of food converted to biomass (Mueller, 1990; Joshi and Mueller, 1996). Recent larval crowding studies on *Drosophila* populations, using a shallow food column, reported an increase in efficiency of crowding-adapted larvae, with respect to the time taken to convert the same volume of food to biomass (Nagarajan et al., 2016; Sarangi et al., 2016).

###### 1.1.2.3. Time stage 3: the end stage of larval crowding

The end stage of crowding can be imagined as the time when the food runs out or the waste concentration becomes unbearably high for the larvae (fig. 1.a). At this stage, a different set of traits would help the focal larva survive, all else being equal. In particular, a greater waste tolerance would allow the focal larva to continue eating at a waste concentration intolerable to other larvae (fig. 1.di). Metabolic waste products such as ammonia, uric acid or urea have been shown to increase in crowded cultures in multiple studies (Botella et al., 1985; Borash et al., 1998; Sarangi, 2018). Furthermore, waste tolerance also evolved in populations adapted to crowding at high food volumes (Shiotsugu et al., 1997; Borash et al., 1998; but see Sarangi, 2018), but not in populations adapted to low food volumes (Nagarajan et al., 2016; Sarangi et al., 2016). In cultures with high food volumes, the food column length is greater (assuming similar container dimensions), and there is substantial food remaining even at the end stage of larval crowding (Bierbaum et al., 1989; Mueller, 1990; Venkitachalam et al., 2023b). Thus, it is likely that, when faced with long food columns, some larvae could feed slowly and survive late into the life of a culture, assuming they can tolerate the increased waste build-up expected due to the numerous larvae feeding and excreting in the vial (Borash et al., 1998; Mueller and Barter, 2015). One study showed that offspring of early eclosing adults from crowding adapted populations had higher larval feeding rate and low waste tolerance, whereas offspring of late eclosing adults had greater waste tolerance along with lower larval feeding rate (Borash et al., 1998). This point further reinforces our assumption that the growth stage and end stage of a crowded culture can be quite different ecologically, and have different underlying fitness functions at different times in the culture.

Finally, the end stage may also be tackled by larvae via a different approach – a reduction in the minimum critical mass required for pupation (Bakker, 1961). The focal larva could survive simply by reducing its requirements and commit to wandering at a smaller size (fig. 1.dii). While primarily useful in conditions where food quantities are fleeting, a reduced minimum critical mass could also potentially reduce some of the duration spent in the increasingly uninhabitable environment at the end stage of the culture. No crowding-adapted population till date has been known to show the evolution of reduced mean minimum critical mass to pupation, although some populations have evolved overall smaller sized adults under uncrowded conditions, which might indicate an evolution of smaller minimum critical mass as well (Sarangi, 2018; Venkitachalam et al., 2023a).

## Methods

### 2.1. Recurrent ‘short-hand’ terms

Surface area: cross-sectional surface area of the cylindrical food column in a vial into which this food is cast in experiments.

Column height: starting food column height in a culture vial.

Food volume: starting food volume of a culture vial.

Eggs: starting egg number in a culture vial.

### 2.2. Units used

While our model is simple enough that any units used are at best vague abstractions of reality, the scale at which we have implemented the simulation attempts to work in units relevant to those used experimentally, and which can be followed intuitively. The units used are:

Dry mass (micrograms – μg): most of the reliable mass estimates on larvae and adult flies are found in units of dry mass (Bakker, 1961, 1969; Santos et al., 1997; Sarangi, 2018; Venkitachalam et al., 2023a). Thus, the starting food culture is in units of μg dry mass. The larval body mass is also measured in units of dry mass (μg).

Time (hours): the measurement of development time (egg to wandering, pupal or adult stage) is typically done in hours (Nagarajan et al., 2016; Sarangi et al., 2016). Thus, time steps in the simulation framework are on the scale of hours. This is also logistically useful, as it requires much shorter computation time than implementing time in a per minute scale. The primary caveat is that feeding rate measurements, which are empirically measured at the scale of minutes (e.g., Sewell et al., 1974), have here been converted to the hourly format. This conversion may not be entirely accurate, due to the presence of inter-feeding bouts (Ruiz-Dubreuil et al., 1996) which may be prevalent over longer feeding periods. In our text, ‘hour’ and ‘time step’ are used interchangeably.

Space (mm): the culture set up in cylindrical dimensions, as well as some larval characteristics in the expanded model, such as digging depth and larval cross section surface area, depend on units of space.

### 2.3. A brief overview of the simulation framework

We simulated cultures of given cylindrical dimensions in which ‘larvae’ started as eggs, hatched and then fed until hitting a defined limit. If uncrowded, the larvae stopped feeding after achieving an optimum size and committed to the ‘wandering’ stage. Under crowded conditions, when faced with severe resource limitation or when facing a toxic environment due to high waste build up, the larvae needed to cross their respective minimum critical mass to pupation in order to commit to wandering. Larvae unable to cross this threshold when the environment was no longer habitable ended up ‘dying’.

We started with a ‘base’ model (section 2.4), onto which we subsequently overlaid more nuanced details of larval competition under the ‘expanded’ model (section 2.5).

### 2.4. The ‘Base’ model of larval competition

#### 2.4.1 Simulating uncrowded conditions

The initial model framework attempted to capture the overall patterns of time to reach wandering stage and dry mass achieved at wandering stage, seen in earlier empirical studies in uncrowded cultures at 25°C under constant light conditions and 60-80% relative humidity (Santos et al., 1997). We describe ‘uncrowded’ as a starting egg number of around 70, in a total food volume of 6 mL (400 mm^2^ surface area, 15 mm food column height). We assumed any egg number below 70 in 6 mL of food would give the same culture outcomes in terms of % survivorship, average mass and average development time. Additionally, we did not incorporate any possibility of non-zero density-independent mortality.

#### 2.4.2. Culture set up

Empirically, the food in a culture is usually measured in volume (mL, or mm^3^), which are components of space rather than mass. Given that most laboratory culture media use a very specific ingredient list (e.g., Sarangi et al., 2016), it was relatively simple to convert a given volume of food into dry nutrient mass. We did not partition the food further into macro-nutrient mass. The dry nutrient density for the cornmeal-sugar-yeast medium used in our selection experiments was rounded to 160 μg/mm^3^ for the simulations.

We started by setting up the culture in terms of the food column cross-section surface area and height, which were converted to the total food volume (E1). This food volume was subsequently converted to a corresponding dry mass value (E2).

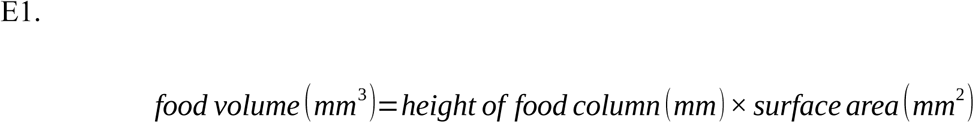

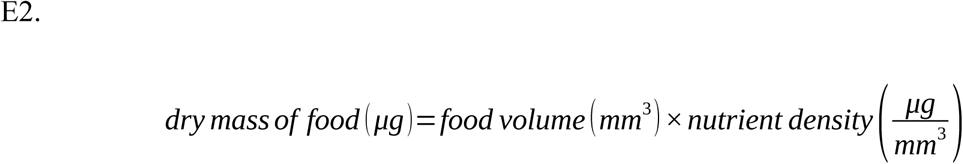

#### 2.4.3. Adding eggs to the culture

A culture with a given food mass was then populated with a specified number of ‘eggs’. The eggs were added simultaneously. These eggs ‘hatched’ into ‘larvae’ which proceeded to feed on the food. Each individual was endowed with a given set of traits (see section 1.1.) when it was first added to the culture. These could be uniform or variable across individuals. When variable, we assumed the traits to be complex enough for their variation to be described by a corresponding normal distribution (but see Borash and Shimada, 2001). Each individual’s trait value was drawn randomly from a normal distribution with given mean and coefficient of variation (see Tables 1-5).

#### 2.4.4. Setting up traits

##### 2.4.4.1. Initial advantage related traits

The details of traits likely to contribute to competitive ability at the initial stage are listed in Table 1. We further calculated egg volume as a cylinder with egg length as height and egg width as diameter. Finally, this was converted to egg dry mass. For this, we assumed that an egg with volume calculated from a mean length of 0.5 mm and mean width of 0.18 mm would have a dry mass of 2.7 μg. This value of mass was obtained from Bakker’s findings (1961), although the association of this mass value with the specific egg volume was arbitrary. However, once the egg volume and egg mass were calibrated in such a way, any combination of egg length and egg width would yield a corresponding egg mass. An individual’s initial larval mass was equal to its egg mass (the mass of the egg shell was not considered – but see Bakker, 1961, 1969).

##### 2.4.4.2. Growth rate related traits

The traits likely to contribute to competitive ability at the larval growth stage are listed in Table 2. Bite rate in uncrowded conditions was assumed to have a positive relationship with larval mass, for a given larva. The relationship was modelled as follows:

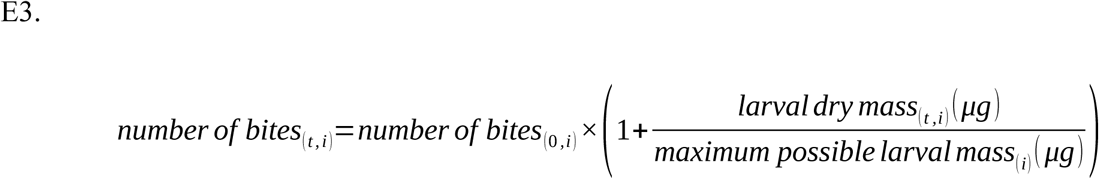

Where *t* = time step (0 = initial time step); *i* = larval identity; and:

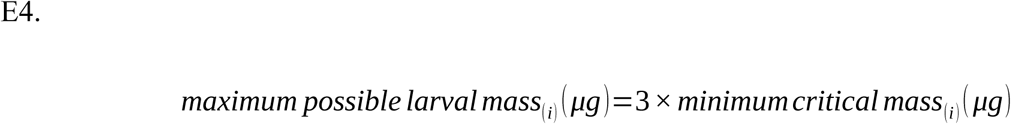

At maximum mass (see Table 3.i), the larva in uncrowded conditions would, thus, have twice the initial bite rate (Santos et al., 1997).

For bite mass, estimates were roughly derived from available larval growth and feeding rate data (Santos et al., 1997), assuming an average food conversion efficiency of 30% (Table 2.iii). We further used extensive mouth part length measurements carried out by Alpatov (1929) to get a general idea of the possible order of magnitude for bite mass, calculated from assuming the bite size as a sphere using a fraction of the mouthpart as the diameter (data not shown). Mouth part size has also been shown to increase over the course of larval growth within the same instar (Robertson, 1963). The association bite mass with larval mass was calculated using a bite scaling factor which scaled bite mass of *i*^th^ larva to its body mass at time *t*. The calculations are as follows:

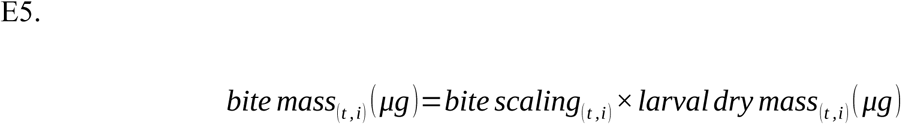

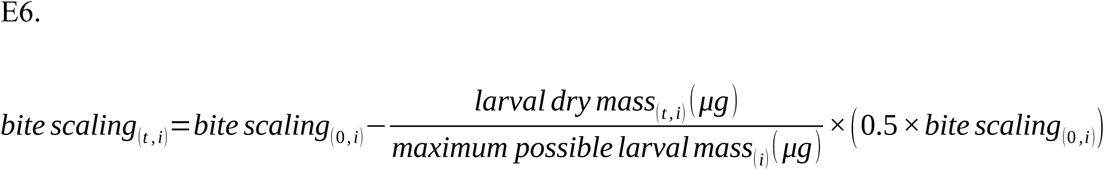

The bite scaling factor had a negative correlation with larval mass. Simulated bite mass increased over the course of larval growth, but the rate of increase of bite mass itself reduced, capping at a minimum rate of zero. Most importantly, we did not consider any variation among larvae in bite scaling (Table 2.ii).

We relied on estimates by de Miranda and Eggleston (1988) to derive values of food to biomass conversion efficiency in *Drosophila* larvae. There were two relevant points in their study – the average efficiency values lay between 20-40% of food eaten depending on the strain, and there was a large amount of variance in estimating efficiency across the time of larval sampling (de Miranda and Eggleston, 1988). Another study by Bakker on multiple populations of *D. melanogaster* found the larval food conversion efficiency values to lie between 20-25% (Bakker, 1969). We accounted for both the mean efficiency and the temporal variation in efficiency per larva (Table 2.iii, iv).

We were unable to find many data regarding the values of time taken to convert food to biomass. However, the study by de Miranda and Eggleston (1988) found the time window to lie within the bounds of 60 minutes. Furthermore, a study on crowding-adapted populations measured food passage time through the gut in the order of minutes (Joshi and Mueller, 1996). Thus, we did not incorporate this facet of larval competitive ability in the current study.

##### 2.4.4.3. End stage related traits

The traits likely contributing to competitive ability towards the end stage of a crowded culture are listed in Table 3.

The waste tolerance threshold per larva was calculated as follows:

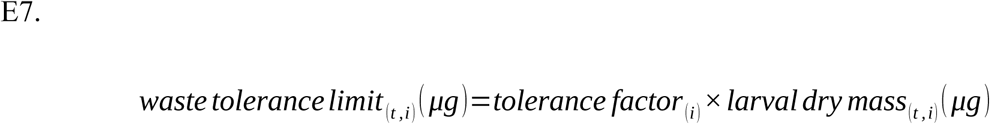

#### 2.4.5. Compiling traits and behaviours

The above factors constituted the larval characteristics in the ‘base’ model, which attempts to capture the understanding of *Drosophila* larval competition as it was around the end of the twentieth century. Each of the traits was incorporated in the process of larval feeding in uncrowded conditions, as well as in various degrees of crowding.

##### 2.4.5.1. Egg hatching

Each added egg started with a counter for its hatching time (Table 1), which reduced by 1 per elapsed time step. After the counter reached 0, the larva hatched at the mass determined by the egg mass (Table 1), and began feeding.

##### 2.4.5.2. Feeding and growth

Every time step *t*, each larva *i* fed as follows:

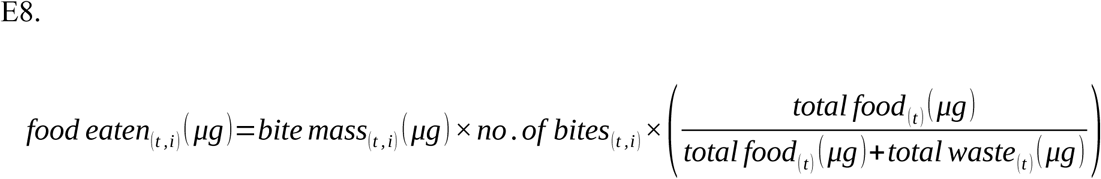

In the base model, growth was simply a property of multiplying food eaten by conversion efficiency, whose value was drawn randomly from a normal distribution every time step. This distribution of possible values had a mean equal to the larval mean efficiency value (Table 2.iii) and coefficient of variation equal to the mean value in Table 2.iv.

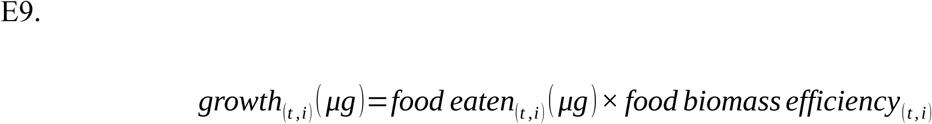

Once the *i*^th^ larva at time *t* crossed its minimum critical mass, it could feed for 36 hours or until it reached its maximum mass, depending on whichever was achieved first. The time limit was derived roughly from the post-critical feeding period seen by Santos et al. (1997), and the knowledge that larvae have a fixed feeding period after achieving their minimum critical mass to pupation (Robertson, 1963; also discussed in Burnet et al., 1977; Prasad and Joshi, 2003). A contrasting result which we did not consider in the current study, was the possible elongation of the third instar in the ‘larval stop’ phenomenon (Ménsua and Moya, 1983; González-Candelas et al., 1990).

Simulated larval growth profiles in an uncrowded condition are shown in figure 2.a, and the outcomes in terms of development time and dry mass at wandering can be found in figure 2.b-c.

**Figure 2.**
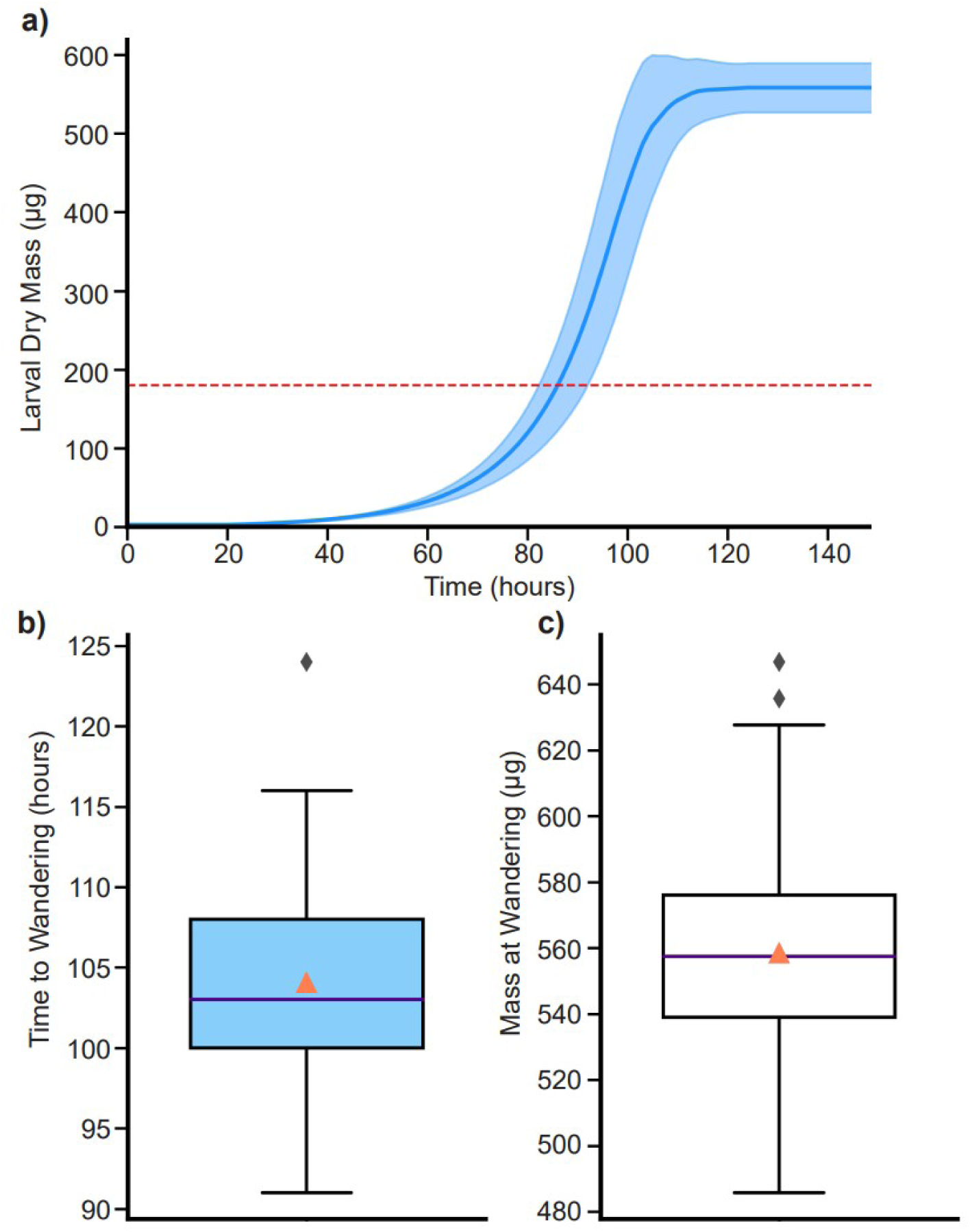
Base model (waste absent), uncrowded: 70 eggs, 6 mL food cast in 400 mm ^2^ surface area and 15 mm height. a) Growth profile for 70 individuals. The solid blue line denotes the mean larval dry mass. The shaded region denotes standard deviation in larval dry mass. The dashed red line denotes the mean minimum critical mass to pupation (180 μg). Once the larvae commit to wandering, their mass remains unchanged. b) Time to wandering distribution. The line within the box denotes the median. The orange triangle denotes the mean. c) Mass at wandering distribution. The line within the box denotes the median. The orange triangle denotes the mean.

In the simulations, if food ran out before a larva completed feeding, there could be two possibilities. If the larva had crossed the minimum critical mass required for pupation, it committed to wandering at the moment of food cessation. However, if the larva had not reached its required minimum critical mass, it died at the moment of food cessation.

##### 2.4.5.3. Waste generation, larval consequences and tolerance

Each feeding larva excreted 2.5% of the total food eaten at the previous time step as metabolic waste (the value used was arbitrary). This waste was representative of, but not limited to, known excretory materials such as uric acid, urea and ammonia, and assumed to be potentially toxic (Botella et al., 1985; Borash et al., 1998; Sarangi, 2018). The waste was excreted by the larva 1 hour after the corresponding food had been eaten (time lag approximated from de Miranda and Eggleston (1988)).

The waste ingested per larva per time step was calculated as follows:

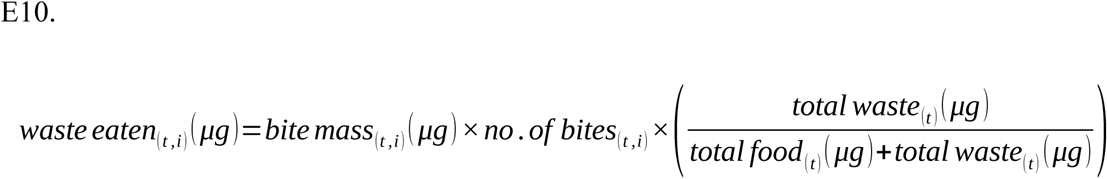

This ingested waste was added to the total pool of waste previously ingested by the larva. If the total waste ingested by a larva crossed its current waste tolerance threshold, then the outcome was similar to food running out. If the larva had crossed its minimum critical mass to pupation, it committed to wandering; if not, it died.

Furthermore, below the waste tolerance threshold, as long as the total waste ingested by a larva was non-zero, it also reduced the bite rate of the larva according to the following non-linear formulation:

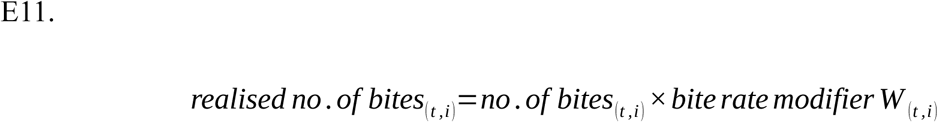

Where,

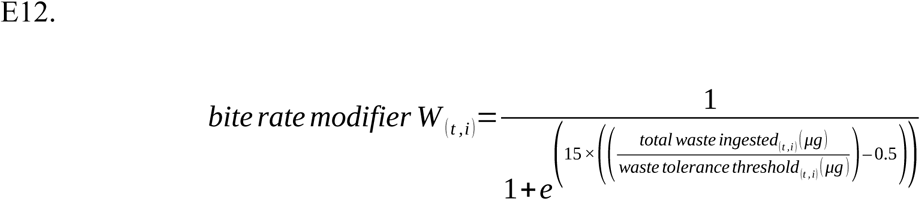

Note: we capped the lowest possible value of the waste ingestion modifier to 0.01, in order to prevent bite rate values from going to 0 and stalling the simulation.

Under the mechanism adopted in this simulation, larvae under uncrowded situations faced very little increase in waste across their growth period, and thus their waste ingested and the associated bite rate modifier values showed little change, leading to very similar uncrowded growth profiles (fig. S3) compared to the base model without waste (fig. 1). In crowded cultures, as food ran out and waste concentration increased, greater waste ingestion caused the bite rate modifier (Eq. 12) to reduce rapidly. Those larvae which had greater waste tolerance reached the inflection point of bite rate reduction at a greater waste concentration.

#### 2.4.6. Base model – introducing larval crowding

Reducing the food volume and/or increasing egg number introduced larval crowding in the simple base model. The results for these with and without the incorporation of waste are given in section 3.1.2.

While our culture set-up method allowed us to vary food volumes in different cylindrical dimensions, the base model described above did not yield differences in cultures with the same egg number and food volume cast into different cylindrical dimensions (detailed in section 3.2 and fig. S4, S5). The base model was thus insufficient in capturing details of recent empirical findings on the effects of egg number, surface area and food column height (Venkitachalam et al., 2023b). For these, we expanded the base model to include mechanisms that could affect crowding along the axes of changing food column cross-section surface area or height, respectively.

### 2.5. The ‘Expanded’ model of larval competition

Recent findings on the importance of feeding band density, as well as the existence of some form of diffusion of metabolic waste into the food column, were incorporated to construct the ‘expanded’ model. We put the disclaimer that the mechanisms for these recently discovered phenomena are not known in detail, and we have put our best speculations into the model, based largely on personal observations, speculations from earlier papers, or logical assumptions about the overall process. These have been classified according to mechanisms and the larval traits involved in interfacing with those mechanisms.

#### 2.5.1. Mechanism: how does food column cross-section surface area affect the outcomes of crowding?

In all likelihood, this is primarily a function of competition for limited space. One of the clearest lines of evidence for this lies in cultures with relatively small surface area and long food columns, wherein much of the food at the bottom is left untouched, presumably due to lack of larval access to food at the bottom of the vial. Moreover, our personal observation as well as those of some other authors (Gilpin, 1974; Botella et al., 1985; Moya and Castro, 1986; Gregg et al., 1990; Chippindale et al., 1994; Mueller and Barter, 2015) have been that crowded larvae tend to feed in packed clusters rather than spread out over the complete surface of the food in contact with the air, even in cultures within very narrow vials. This is presumably done for more efficient feeding by breaking down the food, either physically through mouthparts (Burnet et al., 1977) or through enzymes such as amylase, resulting in easier consumption (Gregg et al., 1990; Sakaguchi and Suzuki, 2013). In conditions of food shortage, this could be particularly important, as easier consumption of food could possibly lead to faster growth.

Thus, the simulated mechanism for limiting surface area was implemented by considering a) difference between total surface area of a vial vs. the effectively available cross-section surface area of the food, as limited by the larval clustering and breaking down of food; and b) difference between effectively available surface area of the food and the total space taken up by the larvae. At the start, each culture had a small fraction of the total vial cross-section surface area as the effectively available surface area. This was meant to represent some initial points which the larvae would start feeding at, which could be chosen either because the food at those points was softer or more fractured, or just arbitrarily as an initial feeding point.

Moreover, we introduced another larval measurement – its cross-section surface area. The initial larval cross-section surface area translated directly from the cross-section surface area of the egg (Table 1.c). As a larva started feeding and growing, its own cross-section surface area also increased in proportion to its mass. This was calculated by assuming (arbitrarily) that a larva with a minimum critical mass of 180 μg would have a cross-section surface area of around 1.54 mm^2^ when at its maximum mass. The larval surface area was calculated from a cross-section diameter of 1.4 mm at maximum mass, measured by Kaznowski et al. (1985). When the larvae, each with their own cross-section surface area, fed in a column of food, they occupied a proportion of the effectively available surface area. The occupancy of the effective surface area was calculated by dividing the total larval cross-section surface area in a culture, by the effective surface area of the vial. In case of a large number of larvae present in a limited vial surface area, as was expected to occur in crowded scenarios, the effective surface area occupancy was likely to be >=1 (fig. 4.ci, cii). Meanwhile, larval feeding activity would also be expected to further loosen and liquefy the surrounding substrate over time, which we represented by the effective available surface area increasing every time step, the rate of which was dependent on whether or not the summed larval surface area completely filled up the available surface area.

We defined a ‘crowded’ situation as one where the effectively available surface area was at least completely occupied by feeding larvae. Besides the clustering feeding behaviour observed, a recent study revealed an additional plastic behaviour that exists for crowded vs. uncrowded situations in larvae. A set of crowding adapted populations showed greater larval feeding rate only under crowded conditions (Sarangi, 2018), but not when assayed singly (or in small groups) on a petri plate (Sarangi et al., 2016; Sarangi, 2018). We thus implemented a plastic increase in feeding rate in the simulation framework, triggered by crowding of the surface area. This was done through the incorporation of two additional traits in the larvae – crowding threshold and peak bite rate modifier (Table 4). We assumed that larvae could feed faster if they detected crowded conditions, as represented by each larva’s crowding threshold. A larva with a lower threshold could potentially detect the presence of crowding at a lower surface occupancy than other larvae, and begin feeding faster a few hours before the other larvae detected crowding and increased their own bite rate. The degree to which the feeding could become faster was determined by the peak bite rate modifier (Table 4). The realised bite rate of a given larva per time step was its bite rate modifier (suffixed *C* i.e., under crowding) multiplied by its bite rate under uncrowded conditions. This bite rate modifier was calculated as follows for each larva *i* at time step *t*:

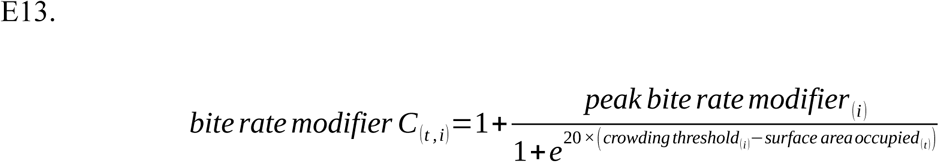

**Table 4.** Traits that may be important to competitive ability under crowding due to surface area limitation.

| **Trait** | **Significance** | **Notes** | **Mean** | **c.o.v.** | **Source(s)** |
| --- | --- | --- | --- | --- | --- |
| **i. Crowding threshold** | Determined the occupancy of the effective surface area at which the bite rate modifier peaked in value. | This was the threshold of space limitation at which a larva detected ‘crowding’. | 1.05 | 5% | Arbitrary |
| **ii. Peak bite rate modifier** | Determined the peak bite rate at the crowding threshold. See equation E13. | This represented the amelioration of detrimental effects of crowding for limited space. | 1 | 5% | Sarangi, 2018 |

Moreover, there was a detrimental effect of crowding for limited space, represented by a loss of food eaten per time step. Our imagined mechanism was the following:

As the larvae crowded around limited cluster spots, they collided against and displaced one another (also speculated by Bakker, 1961; Gilpin, 1974), and not all bite attempts led to food being consumed. We further assumed that larger larvae were better able to embed themselves into the food than smaller larvae (also speculated by Gilpin, 1974), and were thus given less disadvantage in terms of loss of food eaten.

The realised food eaten by a larva per time step was the food eaten under uncrowded conditions multiplied by the food eaten modifier at that time step. The mass percentile of a given larva *i* at time step *t* was calculated for its larval mass as compared to all other larvae at that time step.

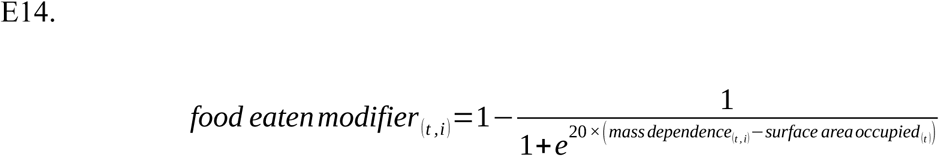

Where:

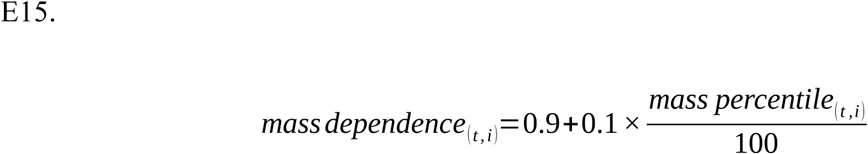

The minimum value of food eaten modifier was set to 0.01 – food eaten per time step could not dip below a hundredth of the uncrowded value. This was done to prevent stalling the simulation.

#### 2.5.2. Mechanism: how does food column height affect the outcomes of crowding?

The presence of long food columns under conditions of crowding has a very consistent outcome – the presence of relatively unused food at the bottom of the vial. This has been discussed in previous crowding-adaptation studies (Bierbaum et al., 1989; Mueller, 1990). Our recent study also showed similar outcomes in conditions with a narrow surface area and a deep food column (Venkitachalam et al., 2023b). Likely due to their need for access to air, larvae tend to feed with their mouths facing downwards, and the posterior spiracles facing the food surface in contact with the air (Sang, 1949; Green et al., 1983; Ruiz-Dubreuil et al., 1996). The larvae can also dig to a certain length in order to procure food (Sang, 1949). When faced with intense crowding, smaller larvae may be unable to dig too deep, as they could subsequently fail to come back up for air and thus drown. Moreover, the larvae excrete the waste material (along with egestion) via the anus (reviewed in Kuraishi et al., 2015) at the top of the food when facing downwards. It has been speculated that under crowded conditions, larger larvae would be able to feed on relatively untainted food whilst polluting the food for smaller larvae closer to the surface of the food (Gilpin, 1974). An earlier study from our group has also given evidence for the presence of larval metabolic waste at the lower half of long food columns after a period of 4-7 days, while the waste at the top saturates around the same period (Sarangi, 2018).

Consequently, we implemented a mechanism for the permeation of waste into the food column over time. The food column was divided into 0.01 mm thin uniform ‘layers’ – each layer having some uniform capacity to hold a certain dry mass of waste in it (arbitrarily, 4.5% waste concentration per layer). As the larvae fed and excreted, the waste first entered the topmost food layer. With waste build up, this layer reached its waste saturation limit at some time point. Subsequently excreted waste went down to the layer below the top layer. Once that got saturated, the process repeated and waste filled up the next layer, with layers continuing to get saturated with waste in a top-down manner (fig. 4a, b). If all the layers were saturated with waste, the new waste was deposited only on the top layer, with the assumption that the rest of the layers were unable to hold any more waste (fig. 4a).

In addition to dry mass and cross-section surface area, we also assigned a length (mouth to anus) characteristic to larvae. Larval length helped determine how deep a larva could dig and access food. The initial length of a larva translated directly from its egg length (Table 1b). As done for larval cross-section surface area, a larva having minimum critical mass of 180 μg was arbitrarily associated with a length of 4.6 mm at maximum mass. This maximum length measurement was obtained from Alpatov, (1929), as well as Kaznowski et al. (1985) (also verified by Bakker, 1959).

Furthermore, each simulated larva was assigned a trait, digging length multiplier, which could vary across individuals (Table 5). This value, multiplied by larval length at a given time step, determined the maximum digging distance for a given larva (fig. 4bi, bii). At any given time, an actively feeding larva could access certain food layers containing 0-4.5% waste concentration. It was assumed that a larva could access food layers from near its mouth position, down to its maximum digging length. The furthest food layer accessible away from the top layer was rounded up from the digging length of the larva. The nearest food layer accessible from the top was rounded down from the larval length. A larva only faced the waste concentration available in accessible food layers. For example, if only layers 1-3 from the top were saturated with waste, while a given larva was feeding at the layers 4-6 from the top, it experienced no waste. If the total digging distance of a larva were to exceed the total number of food layers, it then experienced the entirety of waste in the totality of food. Empirically, this is likely to occur in scenarios where the food column length is shallow or the food gets consumed, resulting in a shallow food column length (fig. 4a).

**Table 5.** Traits that may be important to competitive ability under crowding with long food columns.

| Trait | Significance | Notes | Mean | c.o.v. | Source(s) |
| --- | --- | --- | --- | --- | --- |
| Digging length multiplier | Determined the depth to which a larva could dig, as a proportion of its own body length. | Digging depth = current larval length × digging multiplier | 1.5 | 5% | Inspired by discussion in Sang, 1949 |

### 2.6. Total treatments used – replication of experimental design

In order to capture the full scope of experimental results seen, we attempted to replicate the design used in our recent study (Venkitachalam et al., 2023b). This design consisted of a 3×3×3 factorial design with egg number, food column height and vial surface area as factors, respectively. The egg numbers used were 200, 400, 600; food column height levels were 5, 10 and 15 mm respectively; surface area levels were 200, 400 and 600 mm^2^, respectively. The entire setup was run for 30 replicates per treatment combination. An expansion of this framework for predictive purposes is described in section 4.2.

### 2.7. Note on absence of Statistics

Following the reasoning used by White et al. (2014), we have refrained from performing statistical analyses on our simulation results, instead focusing more on the patterns of differences seen among simulation runs with varying parameter values. In brief, given that we can replicate our simulations manifold for cheap, it is inappropriate to use statistical tests designed for comparisons in experiments which are resource intensive and involve limited replication. Most factors used in greatly replicated simulations would be statistically significant simply from having artificially inflated statistical power (White et al., 2014). Moreover, when trying to capture patterns seen in the experimental results with our simulations, we already knew that a subset of parameter combinations mimicking the treatment combinations in Venkitachalam et al. (2023b) would yield very different outcomes, based on prior limited simulation runs. Thus, as White et al. (2014) note, we knew *a priori* that the null hypothesis of no effect of different parameter combinations was false. Therefore, rather than do significance testing, we focused instead on the overall patterns observed in our simulated datasets, and compared them with corresponding patterns in empirical data. We further extrapolated our simulations to make gross predictions of patterns beyond the reference experimental set up used (section 4.2).

### 2.8. Simulation code

All code was written on Python 3 (Van Rossum and Drake, 2009), using the NumPy package (Harris et al., 2020). Plots were created using Matplotlib (Hunter, 2007) and Seaborn (Waskom et al., 2017) packages.

All the code for running the simulation can be found at the following GitHub link

(Note: this is a private repository until publication – please contact authors for access to code if required prior to publication): https://github.com/SrikantVenkitachalam/Single-Generation-Simulation-Framework.git

## Results

### 3.1.1. Base model – uncrowded conditions

The growth profiles over time for 70 individuals in 6 mL food (400 mm^2^ surface area, 15 mm height) are shown in figure 2.a. The corresponding time to wandering and mass at wandering data are shown in figure 2.bi, bii. These closely corresponded to empirical values observed (Santos et al., 1997).

### 3.1.2. Base model – with larval crowding

The results of increasing egg number from 70-700 in 2 mL (400 mm^2^ surface area, 5 mm height) food are given in figures S1, S2. Results shown in fig. S1 were generated using the base model without considering the effects of waste. Results in fig. S2 had the added effect of waste in the base model. These show the changes in egg to wandering survivorship (fig. S1.a, S2.a), as well as distributions of time to wandering (fig. S1.b, S2.b) and mass at wandering, respectively (fig. S1.c, S2.c). While patterns of survivorship and mass at wandering were largely similar with or without the addition of waste, it was in the distributions of time to wandering that the effects of waste came into play (fig. S1.b vs. S2.b). Without waste, the mean development time decreased across increasing egg number, and the variation collapsed (fig. S1.b). When the effects of waste were considered, mean development time increased, along with an increase in the variation around the mean (fig. S2.b).

### 3.2. A factorial framework for larval crowding

In addition to simpler consideration of changes only in egg number in fixed volumes of food (fig. S1, S2), we also tested the base model in the fully factorial framework of egg number, food column cross-section surface area and height.

This was a simulated ‘replica’ of the experiment reported in Venkitachalam et al. (2023b), examining 3 levels of egg numbers (200, 400 and 600 eggs), 3 levels of food column height (∼5, ∼10 and ∼15 mm) and 3 levels of food column cross-section surface area (∼200, ∼400 and ∼600 mm^2^), in a fully factorial design. Within these 27 treatments lay several pairs of the same combination of egg and food volume, cast into different cylindrical dimensions (varying height and surface area). In total, there were 9 such pairs (200, 400, 600 eggs in two variants each of 2-, 3- and 6-mL food, respectively).

We implemented this experimental paradigm in our base simulation framework (with waste) for egg to wandering survivorship (fig. S4) and time to wandering (fig. S5). Cultures with the same food volume are highlighted using colour-matched text (e.g., fig. S4.i and S4.ix). In the base model, there were no differences observed for any pair of cultures containing the same volume of food, contrary to the experimental observations (Venkitachalam et al., 2023b).

### 3.3. The Expanded model

The 27-combination simulation of the experiment described by Venkitachalam et al. (2023b) was also run for the expanded model, incorporating waste diffusion, variable digging lengths and crowding for limited realised vial surface area (Methods section 2.5).

With the expanded model, there were major differences under various conditions of crowding compared to the base model (fig. 3 vs. fig. S4), while no qualitative differences were seen under uncrowded conditions (fig. 2 vs. fig. S6). Figure 3 shows the egg to wandering survivorship obtained from the expanded model simulations for the 27 treatments described in section 3.2, with 30 randomly seeded replicates for each. Figure S7 shows the corresponding mean time to wandering data. Some of the salient observations are as follows:

1. On average, survivorship decreased as egg number increased under every food culture (fig. 3.d). This was also observed in empirical data (all empirical comparisons in this list are published in Venkitachalam et al. (2023b)).
2. Survivorship increased with an increase in surface area (fig. 3.c) – also observed in empirical data.
3. Survivorship increased with increase in column height only for the transition from 5mm to 10mm (fig. 3.b). Even in that, there was a very large variance in data. In the base model, increasing height improved survivorship in the exact same way as increasing surface area did (fig. S4.b vs. S4.c). In empirical data, height increase also resulted in increasing survivorship, but to a very low degree compared to egg number and surface area. This result was thus close to what was seen in the expanded model. Furthermore, the more prominent change was from 5 mm to 10 mm height, similar to our model, although in the empirical data there was also a small but significant change observed from 10 mm to 15 mm, unlike in the simulation (fig. 3.b).
4. For each pair of cultures containing the same egg number and food volume, survivorship was higher in cultures with greater surface area and shallow depth of food, as compared to narrow surface area and long food column (fig. 3.a. i vs. ix, 3.a. iv vs. viii, 3.a. ii vs. vi). This was consistent with the empirical data. This pattern was not captured by the base model (fig. S4).
5. Mean time to wandering increased with increasing egg number, and decreased with increasing vial surface area (fig. S7.d, S7.c, respectively). This was consistent with empirical results.
6. There was a non-linear change in mean time to wandering along the food column height axis (fig. S7.b). Time to wandering increased from 5 mm to 10 mm height. However, there was a slight decrease in the overall time to wandering from 10 mm to 15 mm height. This was unlike the empirical results, wherein the development time increased almost monotonically, albeit to a very small degree, from 5 mm to 15 mm height. We speculate about possible explanations for this discrepancy in discussion section 4.3.1.
7. For pairwise simulated comparisons of the same combination of egg number and food volume (fig. S7.a. i vs. ix, S7.a. ii vs. vi, S7.a. iv vs. viii), mean time to wandering was higher across all egg numbers in the deeper food column with narrower surface area in 2 mL and 3 mL food. In case of 6 mL, the culture cast in the deeper column with the narrower surface area had similar time to wandering at 200 eggs and 400 eggs, but had lower time to wandering at 600 eggs than its culture counterpart cast into a shallower and wider cylinder. In the empirical results, the egg to adult development time was greater in the deeper versions of each food volume only at the higher egg numbers, and not at 200 eggs. This pattern was particularly exaggerated at 3 mL food.
8. In simulation results, within the deeper versions of 2 mL and 3 mL food, there was little difference in time to wandering across increasing egg numbers (fig. S7.a.iv and S7.a.i, respectively). In the experiment, there was a clear increase in development time across increasing egg numbers in these cultures – particularly in 3 mL food.

**Figure 3.**
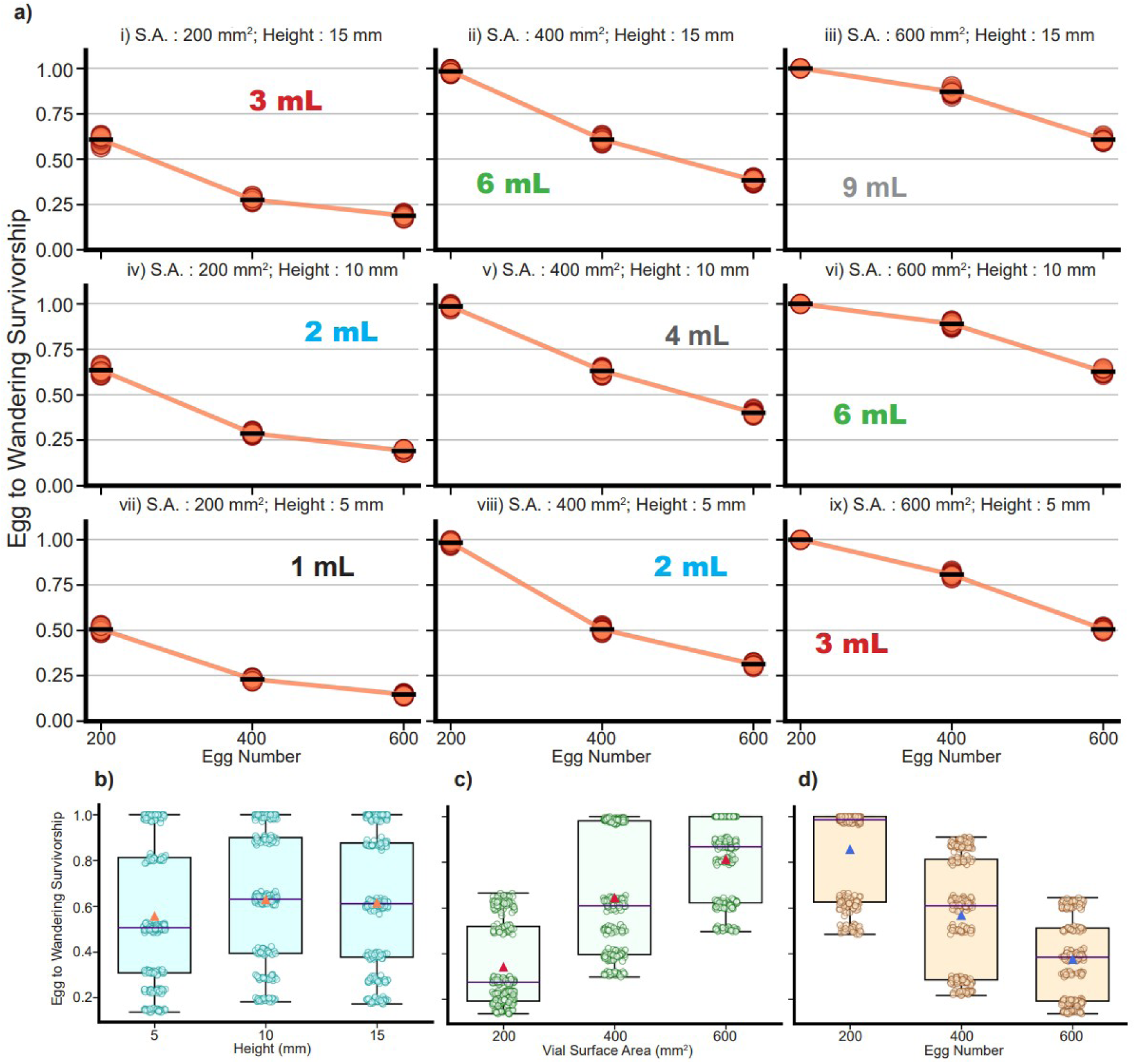
Egg to wandering survivorship in a large factorial simulated experiment (see Methods). S.A.: Cross-section surface area of the food column; Height: starting height of the food column. a) Survivorship of 30 replicates for each of the 27 simulated treatments. The ‘-’ symbol in black represents the overall mean for 30 replicates for each treatment. The food volume is also highlighted for each surface area and column height combination. b) Survivorship values for the three levels of food column height pooling data across all other factors. The indigo line within the box represents median, the orange triangle represents the mean. c) Survivorship values for the three levels of surface area of the food column, pooling data across all other factors. The indigo line within the box represents median, the red triangle represents the mean. d) Survivorship values for the three levels of starting egg numbers, pooling data across all other factors. The indigo line within the box represents median, the blue triangle represents the mean.

As can be seen from the results, there were several differences in survivorship and time to wandering when the same number of eggs were added to the same volume of food cast into cylinders of different dimensions. We further elucidated the differences between the most extreme such pair of the crowded cultures – 600 eggs in 3 mL food – through differences seen in waste permeation, food depletion, digging depth distribution and crowding of realised vial surface area (fig. 4).

**Figure 4.**
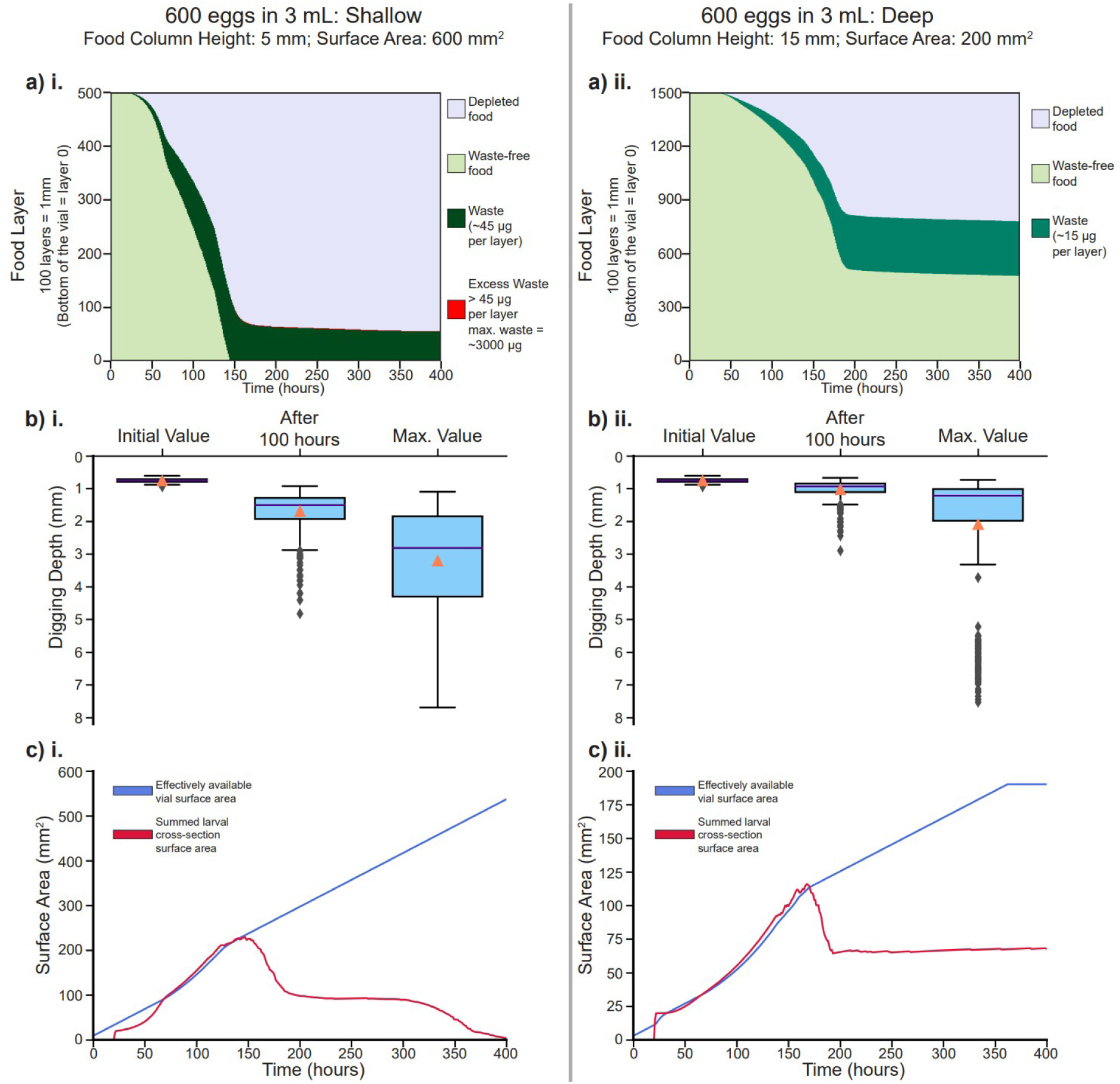
Properties of two simulated cultures with the same egg number and food volume, cast into two different cylindrical dimensions of food. The same individuals are initially seeded in both cultures. The left column denotes properties of the ‘Shallow’ type culture (see figure title); the right column denotes properties of the ‘Deep’ type of culture. a) Waste permeation and food depletion over time in both cultures. Each point on the X-axis denotes a temporal snapshot of the status of the food layers presented in the Y-axis (see key).

i. In the shallow type culture, the food depletes rapidly, until less than 1 mm deep food remains after 150 hours. At this point, all layers of food are saturated with waste, and freshly excreted waste pools at the top food layer. This excess waste can be seen as a thin red line at the interface of depleted food layers and waste saturated food.
ii. In the deep type culture, food depletes until about 175 hours, after which the depletion starts plateauing. Almost half the food layers remain untouched by waste. Over half the food remains in the culture even after 400 hours, and little changes afterwards (data not shown). b) Digging depth of larvae – showing the initial digging depth of freshly hatched larvae, at 100 hours from simulation start, and the maximum digging depth achieved by each larva, just before the point of wandering or death. While the initial distributions are the same in both cultures, the mean, median and quartile range of the shallow culture (b.i.) exceeds that of the deep culture (b.ii.) from 100 hours onwards, until the maximum. In case of the deep culture, only a few larvae achieve high values of digging depth, exceeding 5 mm, signified by outliers in b.ii., while most larvae remain small and dig to depths within 3 mm. In the shallow type culture, the mean digging depth at the max. point is itself at 3 mm, and many more larvae feed upto depths >3 mm. c) The crowding dynamics of both types of cultures, signified by the value of summed larval cross section surface area (red) in relation to the expanding values of the effectively available vial surface area (blue). In both cultures, the summed larval cross section is initially zero when larvae are in their egg stage, and the value becomes >0 after hatching. We define crowding as the time when the summed larval cross-section surface area exceeds the effectively available vial surface area. The shallow type of culture (c.i.) has a much smaller window of time (from 75 hours to 150 hours) when it undergoes crowding, compared to the deep type culture (c.ii.), wherein crowding starts right from the moment of hatching at around 20 hours, and lasts until around 175 hours.

In our empirical study, major insight was gained by considering the outcome of crowding in terms of larval feeding band density (or effective density) instead of the total (eggs/food) density (Venkitachalam et al., 2023b). We defined the feeding band as the shallow volume of food in contact with air, to which larval feeding remains restricted. We had considered the top food volume of up to 6 mm depth as the feeding band (this value was reduced to the total volume of food if starting column height was lower than 6 mm).

We carried out the same exercise with our simulation results replicating the experimental design (fig. 3, fig. S7). We tested for similarities in the overall results with respect to cultures having the same total density across different effective densities. We considered the most crowded case of 200 eggs/mL total density. There were 5 cultures in total – 200 eggs in 1 mL, 400 eggs in 2 mL cast into deep or shallow cylinders respectively, and 600 eggs in 3 mL cast into deep or shallow cylinders respectively (both 2 mL and 3 mL cultures also had corresponding changes in cross-section surface area along with changes in height).

The survivorship and mean time to wandering of these simulated cultures are plotted in fig. 5. Similar to empirical results, there was no discernible change in both fitness related traits when total and effective density were the same (fig. 5.a.i., fig. 5.a.ii), accomplished by keeping the height constant and changing food volume via surface area, regardless of the egg and food combination used. In case the effective density exceeded the total density, accomplished by changing food column height while keeping vial surface area constant, survivorship decreased monotonically with increasing egg number (fig. 5.a.i). This was seen in empirical results as well (Venkitachalam et al., 2023b). While the increase in mean development time with increase in egg number was monotonic in case of experimental results, our simulations differed at least partly. While there was a relatively large increase in average time to wandering from 200 eggs 1 mL to 400 eggs 2 mL, the overall mean time to wandering fell somewhat from 400 eggs 2 mL to 600 eggs 3 mL (fig. 5.a.ii). We address this discrepancy in section 4.3.1.

**Figure 5.**
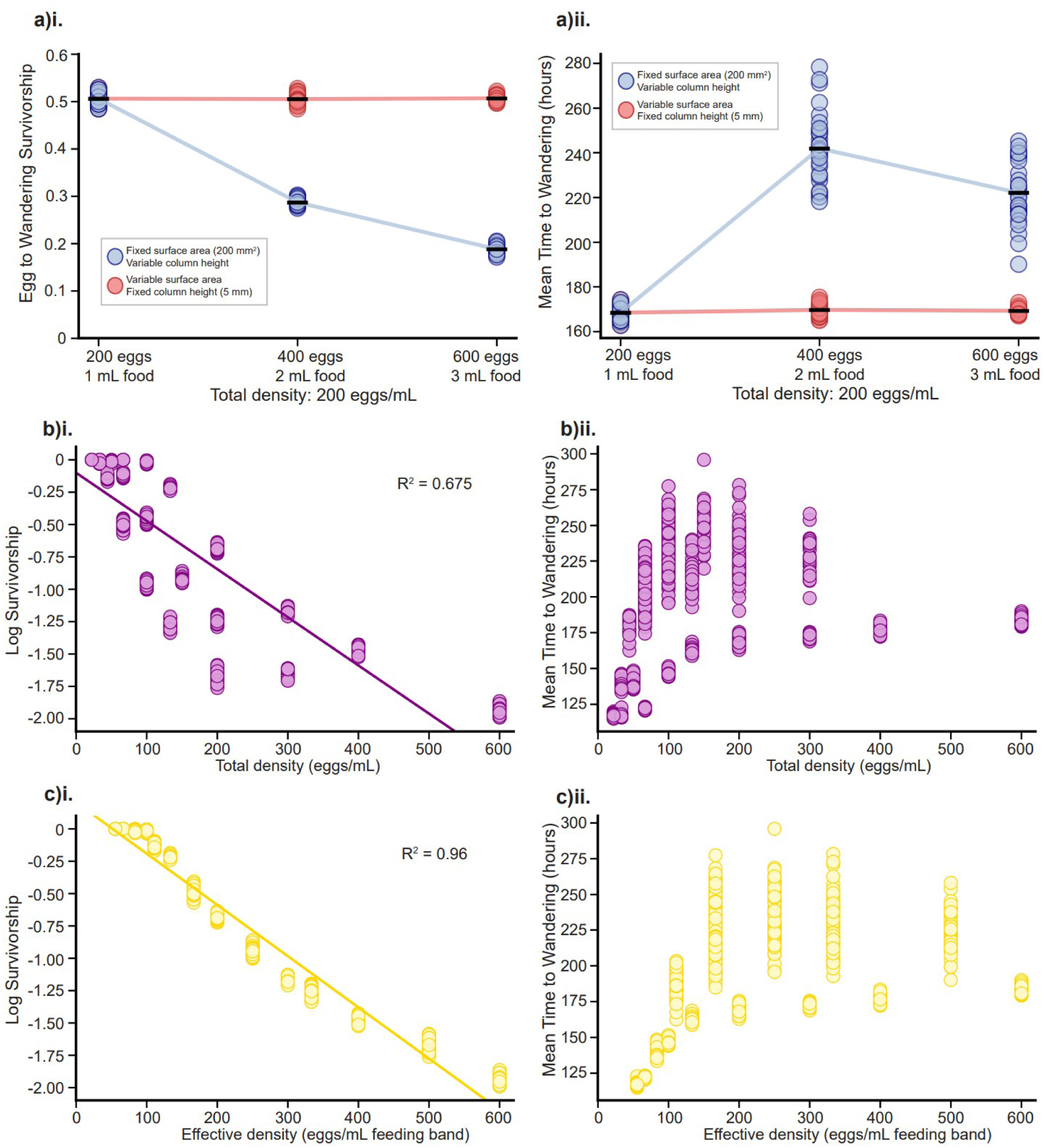
a) i) Survivorship and ii) Mean time to wandering for each simulated culture having 200 eggs/mL total density. The ‘-’ symbol in black represents the overall mean for 30 replicates for each culture type. Plots b.i. and c.i. show the linear regression of log survivorship (pooled data from simulated experiment seen in fig. 3), as predicted by total density and effective density, respectively. b.ii. and c.ii. show the scatter of mean time to wandering of the pooled data, vs. total density and effective density, respectively.

Finally, we pooled the entire dataset with respect to survivorship and carried out a simple linear regression using either total density or effective density (i.e., feeding band density) as predictor variables. The survivorship data were log transformed for linearisation, as the drop in survivorship from low to high density was non-linear. This was unlike the empirical results wherein there was non-zero mortality even at the lowest densities used, leading to an overall linear relationship between total density and survivorship (Venkitachalam et al., 2023b). In the experimental paradigm used, the decreased survivorship at the relatively lower (but not uncrowded) densities was likely a combined effect of density-dependent and density-independent mortality, the latter not having been considered in our simulation framework.

Linear regression for survivorship as the dependent variable in the simulated data showed patterns very similar to the empirical results – effective density could explain far more variation for log survivorship than total density could. Total density as a predictor variable had an *R^2^* of 0.68 (fig. 5.b.i), while effective density as a predictor variable had an *R^2^* of 0.96 (fig. 5.c.i)

For time to wandering, however, a simple linear regression could not be carried out. Unlike in the experimental data, the simulated development time data were very clearly showing two different clusters changing differently with total or effective density (fig. 5.b.ii and 5.c.ii). The part of the data that had different total and effective densities, in case of survivorship, came to lie more closely on a line predicted from data that had the same total and effective densities. This was largely not the case for time to wandering.

## Discussion

### 4.1. Findings from overall results

Similar to earlier models (e.g., de Jong, 1976; Nunney, 1983; see Introduction), the base model in our simulation framework captured Bakker’s (1961) data quite well (fig. S1.a, S1.b). However, we also found that the base model did not account for changes in development time (fig. S1.c), which is an important fitness-related trait affected by larval competition (reviewed in Venkitachalam et al., 2023a). To account for development time variation, we introduced waste into the system, whose ingestion slowed the overall development time of our model larvae, in addition to killing the ones which ingested too much waste without having achieved their respective minimum critical mass (section 2.4.5.3). The resultant simulation better captured patterns seen in experiments done until the end of the twentieth century, in terms of survivorship, development time and dry mass (fig. S2).

We note that Bakker’s (1961) data, quite uniquely in hindsight (given the many exceptions seen in subsequent studies), represented a purely “exploitation” mode of competition among larvae (*sensu* Park, 1954; reviewed in Birch, 1957), wherein differences in feeding rates were enough to predict the competitive outcomes of various strains of larvae. The crowded cultures in that study were implemented by seeding larvae over a thin layer of yeast covering a thick layer of non-nutritive agar (Bakker, 1961). This may have dispersed a majority of the waste away from the top food-containing layer where the larvae had to feed. Consequently, the near-absence of waste would make competition almost purely exploitative.

This reasoning may explain the success of several models, including our base model, in capturing patterns of data seen by Bakker in his studies. The exploitation type of competition was confounded by Bakker with the term “scramble” (Bakker, 1969), which was described originally to mean incomplete success in competition (Nicholson, 1954). In her model, de Jong (1976) described *Drosophila* competition as scramble + exploitative, i.e., the outcome of larval competition would lead to incomplete success for the survivors as a majority would not feed optimally, and even dying larvae would consume non-zero food. This incomplete success would be expected to come about through purely exploitative action – the larvae would not affect directly or indirectly the feeding process of other larvae (de Jong, 1976).

However, once larvae face increasing concentrations of metabolic waste, as is expected under crowded conditions in homogenous food media, competition may no longer be purely exploitative. Excreted larval waste, which potentially affects the survival and/or development of all competing individuals (Botella et al., 1985), adds elements of indirect interference (Park, 1954; Birch, 1957; de Jong, 1976) into the process of competition (also see Weisbrot, 1966; Dawood and Strickberger, 1969). This could differentially affect individuals, and thus add a new dimension to larval competitive ability (Shiotsugu et al., 1997; Borash et al., 1998). We suspect that *Drosophila* larval competition in most laboratory scenarios would be scramble type with components of both exploitation as well as indirect interference.

A reduction in bite rate would be expected to delay overall development time of the larva under any condition where waste is high. This reasoning follows from discussion by Botella et al. (1985), whose study demonstrated higher waste conditions due to crowding leading to elongated egg to adult development times. We also know from previous larval crowding work that average development time tends to increase under crowded conditions (Sang, 1949; González-Candelas et al., 1990; Sarangi, 2018; Venkitachalam et al., 2023a,b). Mechanistically, this would make sense if a larva has to slow down its bite rate in order to optimally detoxify the accumulating waste within its body. To affirm this mechanism, we would need to test the feeding rate of larvae in food laced with waste products. In the current study, we did not include an actual reduction in waste levels within larvae due to detoxification, although this can certainly be implemented in the future. Moreover, this relationship between waste levels and bite rate could work independently of the knowledge that late-eclosing adults in some crowding-adapted populations evolved reduced feeding rate alongside increased waste tolerance (Borash et al., 1998), or that populations adapted to tolerate high levels of metabolic waste products such as urea or ammonia showed the evolution of lower feeding rates compared to unselected control populations (Borash et al., 2000; Bitner et al., 2021).

While introduction of waste in the simulation framework does replicate patterns seen in the experimental data to a certain extent, there is a fundamental aspect missing. Recent studies conducted by us showed that similar densities cast into different dimensions yield very different outcomes of fitness related traits (Sarangi, 2018; Venkitachalam et al., 2023b). This vital but nuanced aspect of larval competition was not captured by the base model even with the implementation of waste (fig. S4, S5).

To address this, we introduced the expanded model, which includes features that could work specifically with vial cross-section surface area and food column height characteristics (section 2.5). Under this updated framework, simulated larvae could dig into the food column while excreting waste which would saturate the top layers of food first, affecting the smaller larvae which are unable to dig too deep. These larvae would also have their own cross-section surface area. Too many larvae in a constricted cylindrical container would see their cumulative cross-section surface area exceed the relatively little accessible vial surface area, leading to competition for limited space. Ultimately, such crowded conditions could cause larvae to ‘miss out’ on food eaten, while also plastically increasing their bite rate as an ameliorative measure.

These incorporations led to the expanded model capturing several facets of empirical data (see Results 3.3.). In particular, simulated patterns of egg to wandering survivorship mimicked the experimental data quite well (fig. 3). Overall patterns seen in survivorship from simulated cultures having the same combination of eggs and food but cast into different cylindrical dimensions were largely preserved (fig. 3). Similar to empirical observations, there was a lot of unused food left at the end of a simulated culture in a deeper, more constricted food column than when the same egg and food combination was present in a shallow but wider cylinder (fig. 4.a.i, 4.a.ii). The simulated food also depleted faster in wider and shallower cultures than in deeper, constricted cultures – this phenomenon yet remains to be studied experimentally. There was also an overall increase in realised bite rate seen in simulated crowded cultures (data not shown), similar to empirical data (Sarangi, 2018). Perhaps more importantly, regression patterns of survivorship as predicted by total density and effective density (fig. 5.b.i, 5.c.i) were quite consistent with empirical observations (Venkitachalam et al., 2023b). For these regressions, any contradictions of simulated data with those drawn from experiments were in the realm of low total densities, likely due to the absence of density-independent mortality in our model. Effective density in the simulations, like in the experiment, predicted egg to wandering survivorship far better than the total density (fig. 5.b.i, 5.c.i).

While some aspects of development time, represented here by time to wandering, were captured by our simulated data, exceptions were also seen (see section 3.3.). Changes in mean time to wandering along the egg number and vial surface area axes were largely similar to experimental data. However, there was a lack of congruence on the food column height axis. Mean time to wandering increased along with an increase from a very shallow food column (same total and effective density) to a moderate food column (total and effective density not similar). This pattern was consistent with experimental data. However, on increasing food column height further (greater dissimilarity between total and effective density) there was a decrease in mean time to wandering in the simulations, unlike the consistent increase seen in experimental data. This pattern of observations has been explored further in the ‘Predictions’ section below. Along with greater food column height increase, there was also a lack of change in survivorship (fig. 3), whereas in experimental systems it has been well established that survivorship improves when culturing the same number of eggs along an increasing food column height (Sarangi, 2018). However, this improvement is far less than the change in survivorship when vial surface area is increased instead of food column height (Sang, 1949; Venkitachalam et al., 2023b).

### 4.2. Predictions

One of the primary goals of simulating a biological system, such as our current undertaking, is to make predictions beyond the domain of the existing empirical work, ultimately guiding the development of better experimental designs.

After having re-created an existing experimental design within our simulation framework, and achieving reasonable (albeit varying) congruence with empirical results, we simulated two major expansions to the existing experimental paradigm.

The first of these is straightforward. There are three easily observable outcomes of implementing larval competition – reduced egg to adult survivorship, increased mean egg to adult development time and reduced mean adult body mass at eclosion, as compared to the uncrowded conditions (Sang, 1949; Ohba, 1961 – as cited by González-Candelas et al., 1990; Sarangi, 2018; Venkitachalam et al., 2023a). Of these, only the first two were studied experimentally in the factorial experimental framework in Venkitachalam et al. (2023b), and subsequently explored via simulations (fig. 3, fig. S7). Thus, we make broad predictions with respect to body mass distributions in various crowded cultures in the experimental design replicated in this study. The simulated mean and variance in dry body mass at wandering have been shown in fig. 6, fig. S9, respectively. At high surface areas, body mass decreases across increasing egg numbers (fig. 6.a.iii, vi, ix), along with an increase in variance (fig. S9.a.iii, vi, ix). At constricted surface areas there is little change in the mean or variance across egg numbers (fig. 6.a.i, iv, vii; fig. S9.a.i, iv, vii). In deeper and constricted food columns, a similar lack of change was seen for simulated mean and variance in time to wandering (fig. S7.a.i, iv, vii; fig. S8.a.i, iv, vii), but not in empirical data (Venkitachalam et al., 2023b). For the same egg number and food volume, a simulated culture cast into a deeper and more constricted column will have, on average, greater mean and variance in dry body mass (fig. 6.a.i vs. fig. 6.a.ix; fig. S9.a.i. vs. fig. S9.a.ix).

**Figure 6.**
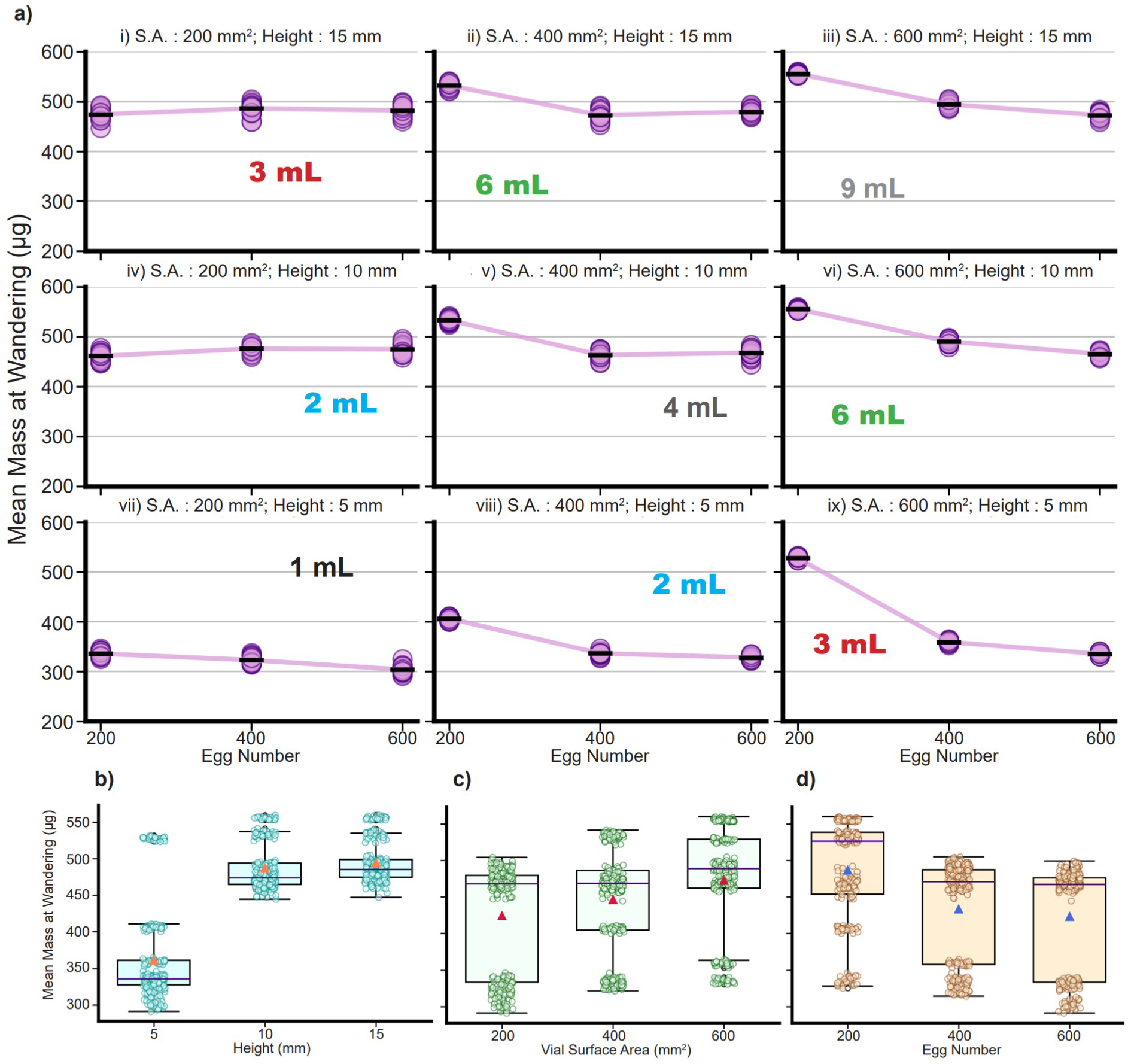
Mean mass at wandering (μg) in a large factorial simulated experiment (see Methods). S.A.: Cross-section surface area of the food column; Height: starting height of the food column. a) Mean mass at wandering of 30 replicates for each of the 27 simulated treatments. The ‘-’ symbol in black represents the overall mean for 30 replicates for each treatment. The food volume is also highlighted for each surface area and column height combination. b) Mean mass at wandering values for the three levels of food column height pooling data across all other factors. The indigo line within the box represents median, the orange triangle represents the mean. c) Mean mass at wandering values for the three levels of surface area of the food column, pooling data across all other factors. The indigo line within the box represents median, the red triangle represents the mean. d) Mean mass at wandering values for the three levels of starting egg numbers, pooling data across all other factors. The indigo line within the box represents median, the blue triangle represents the mean.

The simulations perhaps have a unique situation wherein crowded cultures with lower surface areas have relatively unchanged development time and body mass of survivors across increasing egg numbers. This occurs along with a large reduction in survivorship. In empirical data, this pattern was observed at lower heights, but not at the 15 mm height (Venkitachalam et al., 2023b).

The second major prediction is with respect to the treatment levels used for the same total density. We implemented a total density of 200 eggs/mL across various combinations of eggs and food volume, resulting in a deeper and constricted food column, or a shallow and wide column (as seen in figure. 5.a). The simulated scenarios ranged from 100 eggs in 0.5 mL food, up to 1000 eggs in 5 mL food (fig. 7). We avoided predicting beyond these limits, as other factors such as humidity, or the lack thereof, are likely to become important at very constricted or very wide and shallow food columns. Within the limits seen, there is no change predicted in egg to wandering survivorship or mean time to wandering for increasing surface area (i.e., keeping total density and effective density the same) (fig. 7). When the change in food volume is via column height, egg to wandering survivorship is predicted to decrease consistently with increasing egg number and food volume (fig. 7.a). There is an interesting pattern seen in mean time to wandering – as the food volume increases, the mean time to wandering increases, peaks at 400 eggs in 2 mL food, and reduces to a middle value at higher egg numbers and food volumes, with a slight increase seen with increasing egg and food volumes (fig. 7.b). This is also observed in fig. 5.a.ii, wherein the time to wandering at 400 eggs in 2 mL was greater than in 600 eggs in 3 mL food.

**Figure 7.**
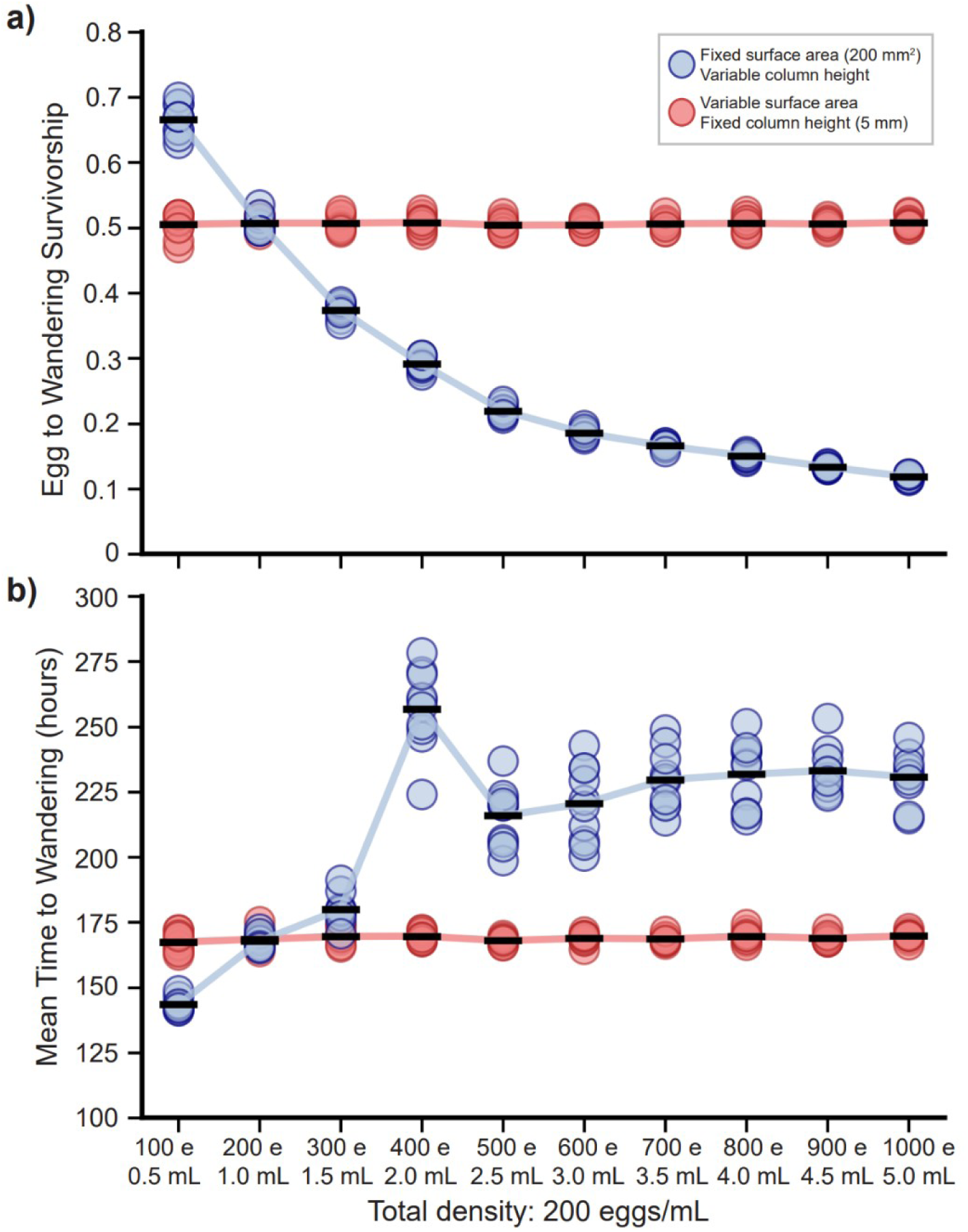
Simulations predicting multiple treatment combinations achieving a total density of 200 eggs/mL. This is an extrapolation of the cultures considered in figure 5.a. The figure shows a) egg to wandering survivorship and b) mean time to wandering for the extended design. The X-axis labels show the egg number (shorthand ‘e’) and food volume (mL) combination used for every treatment pair (see legend). The ‘-’ symbol in black represents the overall mean for 10 replicates for each culture type.

This pattern likely occurs in the simulation because in a culture with 400 eggs constricted into a narrow column with 2 mL food, there is not enough waste accumulated to kill all the larvae as the waste permeates downwards, leading to several waste resistant larvae surviving much later into the life of the culture. Due to the relatively high numbers and lower surface area, there is also enough initial space crowding present such that there would be a further delay in development time. Upon increasing the egg numbers more than 400, the resultant increase in waste levels would kill a majority of the larvae which could have otherwise survived later into the culture if waste concentrations were lower, leading to an overall shorter mean time to wandering (also see Discussion 4.3.1.).

Besides these expansions, however, there is another aspect in which the predictions from the simulation framework can be potentially important – in the elucidation of the mechanisms employed in the process of larval competition.

The formulation of this simulation framework compelled us to peer inside the black box of larval competition, and we found the knowledge of mechanisms wanting. While some traits such as bite rates, egg mass, hatching time and minimum critical mass were fairly well studied and defined, most of our ‘self-defined’ traits (Tables 2-4) were deigned to be so because we did not find any relevant studies that had explored the exact mechanisms or measurements of these traits. These are particularly true for traits, as well as phenomena, used in the expanded model (section 2.5; Tables 4, 5). The mechanisms assumed in our simulations should be good starting points for exploration of these hitherto unexplored traits and characteristics relevant to crowded systems, as they yield a sufficient fit to empirical data. Of greatest import are understanding the following:

a. The complete content, and build-up of metabolic and other egested waste products, as well as their effects on larval feeding behaviour in various kinds of crowded cultures. Knowledge of the mean and variance of waste conversion from food would also be useful when comparing uncrowded and crowding-adapted populations.
b. The overall mechanism by which crowding for limited surface area may occur, as well as its effects on larval feeding behaviour.
c. The dynamics of larval digging – the mean and variance in larval length and digging distance achieved under various scenarios of crowding.
d. The overall phenomenon of waste permeation and/or diffusion – does waste predominantly saturate layer by layer, or does it travel down the food column and settle at the bottom? We did try to implement the latter process in our framework, but the patterns in our simulation results (data not shown) did not capture those seen in empirical data.

So far, none of the points listed above have been pursued experimentally in larval crowding systems, to our knowledge.

We think that further additions and modifications to the mechanisms and traits of the current simulation framework would make more sense once some (if not all) of the implemented mechanisms are empirically better understood. We would, however, be remiss not to credit authors who, several decades prior to our work, made observations or speculations regarding most phenomena and behaviours explored in the current study (Sang, 1949; Bakker, 1961; Gilpin, 1974; Botella et al., 1985; Moya and Castro, 1986; Bierbaum et al., 1989; Mueller, 1990; Chippindale et al., 1994; Sokolowski et al., 1997; Sarangi et al., 2016; Sarangi, 2018).

### 4.3. Drawbacks and possible quirks of the simulation framework

#### 4.3.1. Discrepancies with data

Currently, most of the major discrepancies of simulated results with respect to empirical data lie at high levels of crowding coupled with deep food columns. In simulations, there is no change in survivorship for the same egg number in increasing volumes of food beyond a certain height (fig. 3.a, 3.b). Meanwhile, the simulated time to wandering follows a non-linear pattern across increasing food column heights, both with the same egg number (fig. S7.a.i, iv, vii) as well as proportionately increasing egg numbers (fig. 5.a.ii, fig. 7b) – showing the lowest development time at the lowest food volume, then increasing at moderate volume, then coming to a middle value at a higher volume. In experiments, both survivorship and development time increase for the same number of eggs and increasing food column heights (Venkitachalam et al., 2023b).

There are two possibilities here – a) either we failed to capture the effect of food column height on a crowded culture, or b) we captured the overall pattern but could not capture the corresponding levels of height at which the pattern unfolds. These can be addressed through an expanded experimental design – increasing food volume through several levels of column height while keeping the number of eggs the same. If a) is correct, then both survivorship and development time will likely keep increasing with increasing height (or development time may not change beyond a certain height). However, if b) is correct, there should be no change in survivorship and/or a decrease in development time beyond a certain height. The latter would mean that our simulations captured the overall patterns of the effects of changing food column height, but did not match the dimensions of experimental culture vials correctly. For testing congruence of empirical development time with simulated data, an experiment using uniform total density across several levels of increasing egg numbers and food column heights may also suffice.

The lack of congruence with the outcomes of crowding on changing food column height also creates some discrepancies with data from other studies besides the experiment of Venkitachalam et al. (2023b). We know from previous studies (Sarangi, 2018) the survivorship, pre-adult development time and body mass patterns in cultures crowded at 600 eggs in 1.5 mL food (400 mm^2^ vials), 1200 eggs in 3 mL food (400 mm^2^ vials), and 1200 eggs in 6 mL food (350 mm^2^ vials). While the survivorship data is well captured by the simulations at 600 eggs, 1.5 mL food, there is only a slight reduction in survivorship when doubling the egg number and food volume in our simulations (data not shown), whereas in experiments there is >10% reduction in survivorship (Sarangi 2018). More incongruent are the data from 1200 eggs in 6 mL food – while survivorship is the lowest amongst the three types of cultures in our simulations (data not shown), it is similar to survivorship in 600 eggs in 1.5 mL food, in experiments. This clearly shows the importance of the effect of food column height on survivorship, which does not seem to be captured in our simulation framework beyond a point. However, mean time to wandering and mean mass at eclosion patterns in the respective simulated cultures are very much in line with experimental data (data not shown).

These discrepancies likely happen due to the mechanism of waste permeation considered by us. Given a certain number of eggs and a certain vial surface area, the pattern of waste permeation is likely to be very similar across all but the lowest food column heights. This once again highlights the need for future studies on waste permeation mechanisms and its effects on larval feeding.

An important aspect that has been seen in crowded cultures in 400 mm^2^ vials containing food ≥ 2 mL is the distribution of mass at eclosion across the development time window. Adult mass is maximum for the earliest eclosing flies; those flies eclosing at the middle of the distribution have lower mass; and finally, the last eclosing flies have increased mass, similar to the early eclosing flies (Sarangi, 2018; Venkitachalam et al., 2023a). This pattern of experimental data is likely indicative of some detoxification mechanism in late feeding larvae, or some diffusion of waste away from the feeding band – both of which have not been incorporated in the simulations. Thus, in simulated crowded cultures, while the earliest wandering larvae (and hence eclosing adults) do have the maximum mass, all subsequent wanderers are much smaller, with no larva with delayed development time having increased mass (data not shown).

Finally, while there are several aspects of empirical data that have been successfully captured by our work despite the discrepancies, we do not yet know whether our ‘best-guesses’ employed here actually replicate the process of competition, or if they are simply approximating empirical consequences through non-biological causes. In either case, further experimentation using the leads from the current study are likely to yield greater understanding of the ecology and mechanism of larval competition.

#### 4.3.2. Some additional features of the simulation that may have biological relevance

The following features (or quirks) of our simulation may be worth exploring, for they may represent important real-world relationships.

The first is the effect of waste on the outcome of crowding. We have observed that each time any waste is present in the base model, it becomes the predominant factor controlling the outcome of the culture (fig. S2). The larvae with the highest waste tolerance are the only ones that survive when faced with any amount of waste. No other larvae can survive regardless of any other trait advantage they may possess (data not shown). We know that a host of traits besides waste tolerance can evolve under experiments on adaptation to larval crowding (section 1.1). Thus, it is unlikely that waste is either this prominent or ever-present. As a possible answer to this, the incorporation of waste permeation in our simulations delayed the exposure of waste to most larvae, thus painting what might be a realistic picture for the role of waste in larval competition. It may also be possible that, in reality, the effects of waste on larvae are completely different from what was implemented in our simulations. Elucidating this possibility would require further experimentation.

Secondly, in order to get any realistic result, efficiency could not be plastically reduced due to high waste or high crowding, such as was done with bite rate or food eaten (section 2.4.5.3; 2.5.1). Any time efficiency was reduced, the larvae under crowded conditions even in relatively high volumes of food rapidly depleted all remaining food without converting most of it to growth. This led to almost no survivorship with completely depleted food from initially high food column lengths – a situation that has not yet been observed empirically. Thus, while greater efficiency may play a vital role in faster larval growth (section 1.1, fig. 1), and be selected for under certain situations of crowding, it may be unlikely to reduce plastically to large degrees under any crowding stress (assuming this feature of the simulation is realistic).

An additional feature is ‘crowding’ as defined in the expanded model (section 2.5.1), wherein we predicted the potential importance of the space occupied by larvae (measured in terms of cross-section surface area) and its changing dynamics with the overall accessible space in the feeding band. In the simulations, this feature was of central importance in the outcome of a crowded larval culture (fig. 4.ci, cii). A question that arises from this is whether high surface area occupancy is what represents ‘true’ larval crowding – which would mean that a crowded larval culture is only crowded for a period of time. True crowding may not be occurring initially and especially towards the end of a culture, due to increased availability of food surface area to the larvae (fig. 4.c; also discussed in Sokolowski et al., 1997). Depending on the vial surface area and the number of eggs seeded, the dynamics of crowding as defined here may be central to predicting the outcomes of competition among larvae.

### 4.4. Looking ahead

#### 4.4.1. Factors left to incorporate

Before concluding, we wish to note several traits, characteristics and behaviours we omitted from our simulation framework, in order to keep it from becoming more complex than it already was. To start with the most well documented characteristic, we decided to remove larval instars from consideration. While earlier versions of our framework did have an implementation of instars (S. Venkitachalam and A. Joshi, *unpublished*), it was unclear if the simulations benefitted from the added feature. It is known that both feeding rate and bite mass can increase continuously over the period of larval growth (Santos et al., 1997; Robertson, 1963, respectively), rather than in discrete steps corresponding with instars. According to Bakker (1961), instar moulting episodes could possibly be relevant in terms of enforcing time away from feeding that could be detrimental to a larva’s competitive ability (see Green et al., 1983).

Another characteristic that we have not incorporated is the possibility of drowning of larvae under limited surface-area crowding in cultures with deep food columns. Given that there is limited space under such conditions, it is likely that there would be scope for larval collisions (Gilpin, 1974), and deep digging larvae may be blocked from access to air altogether, resulting in them drowning. This may be an important ‘disoperative factor’ (see Bakker, 1961) in reducing survival under high feeding-band density conditions, while drowning resistance could be a trait to counter it.

There is likely great room for exploration of the ecology and evolution of larval digging behaviour under crowded conditions. Recent studies have highlighted the possibility of ‘cooperative digging’ in *Drosophila* larvae (Dombrovski et al., 2017, 2019), and it remains to be seen if such phenomena are observed in our laboratory populations. Additional related questions also arise – whether such digging phenomena can show plasticity under different scenarios of crowding and whether the level of synchronicity in digging behaviour can evolve under selection for high-feeding band density coupled with deep food columns. The potency of enzymes such as salivary amylase secreted by digging larvae which can help break down the food substrate for easier consumption (Gregg et al., 1990; Sakaguchi and Suzuki, 2013), or rate of the physical breakdown of food by larval mouthparts (Burnet et al., 1977), are each potential candidate traits for the action of selection under crowded conditions.

Finally, in shallow food columns with high larval density, food can run out, leading to larval death (Sarangi et al., 2016; Venkitachalam et al., 2023a; see fig. 4a.i). When food runs out, we have caused the simulated larvae to instantly commit to wandering in case they have crossed the threshold minimum critical mass to pupation. However, real-world larvae may instead attempt to seek out food through increased locomotion (Green et al., 1983; but see Ruiz-Dubreuil et al., 1996) and arrest their development for several hours if they run out of food. The latter is the mechanism described for the phenomenon of ‘larval stop’ observed in earlier studies (Ménsua and Moya, 1983), which has not yet been incorporated in our simulation framework. It is additionally worth pointing out that the presence of large amounts of waste (Botella et al., 1985), or high surface area crowding can also possibly slow down development time under crowded conditions and lead to ‘larval stop’ like phenomena.

#### 4.4.2. Future expansions of the model

##### 4.4.2.1. An evolutionary extension of the single generation framework

Understanding the importance of different traits under various scenarios of larval crowding would involve the extension of the current simulation framework into a multigenerational model. This would allow the study of evolutionary consequences of rearing larvae in given culture conditions. In this extended framework, the wandering larvae would end up as adults, mate and produce offspring with some rules of inheritance, perhaps a simple additive model for quantitative traits to begin with. For a given crowding regime, the offspring produced from multiple replicates at a single generation would be mixed and resampled into new cultures subjected to the same crowded regime, forming the next generation. This endeavour is currently underway (see Venkitachalam (2017) for a preliminary version of this model).

##### 4.4.2.2. Traits likely to evolve across different types of cultures

As a first pass to predicting the traits that might be important in different kinds of crowded cultures, we have plotted the trait distributions of surviving and dying individuals (fig. S10) in two types of cultures – sharing the same number of eggs (600) in the same volume of food (3 mL) but cast into different types of cylinders (‘shallow’ or ‘deep’ – see fig. 4, S10). A brief overview of our findings is as follows:

###### Traits conferring initial advantage

There were no discernible differences between surviving and dying individuals in terms of their hatching times, in either shallow or deep cultures (fig. S10.1.). Experimentally, reduced hatching time has evolved in some crowding adapted populations, but the extent of differences was less than an hour (Venkitachalam et al., 2022). It is likely that this is not observed in simulations due to the 1-hour minimum resolution used in our simulations and low between-larva variation in hatching time (Table 1.a)

Large differences were observed between surviving and dying individuals in terms of both their egg lengths and egg widths, with survivors having larger eggs on average (fig. S10.2; S10.3). This difference was more exaggerated in case of the deep-type culture (fig. S10.2.aii, bii; S10.3.aii, bii), with most of the early surviving individuals having greater egg lengths and widths (fig. S10.2.cii; S10.3.cii). In the simulation, it is likely that an initial mass advantage is critical to surviving and feeding in the more intensely crowded ‘deep’ type of culture, wherein hatched eggs have to immediately face crowded conditions. Interestingly, a recent study by us showed that greater egg length and width evolved in three different sets of crowding-adapted populations compared to low density reared controls, with the largest eggs being found in the cultures adapted to high effective densities with deep food columns (Venkitachalam et al., 2022).

###### Traits conferring growth rate advantage

Initial bite rate was higher among survivors of both types of simulated cultures (fig. S10.4). However, the extent of difference between bite rate of surviving and dying individuals was greater in the shallow type culture (fig. S10.4.a.i). This may be due to the simulated shallow type cultures having a greater scope for larval growth in the initial period of absence of crowded conditions (fig. 4.c.i). In selection experiments, populations reared in crowded conditions similar to the shallow type cultures did not evolve greater feeding rate than controls when assayed singly (Nagarajan et al., 2016; Sarangi et al., 2016). In contrast, survivors of the shallow simulated cultures would be expected to evolve higher bite rate, which would be visible regardless of culture conditions.

A similar pattern as initial bite rate was also seen for mean efficiency in simulations (fig. S10.7.). However, it is unknown if populations evolved in cultures resembling the shallow type have greater food to biomass conversion efficiency. Populations adapted to cultures similar to the deep type evolved lower efficiency than the controls, which may have been due to trade-offs between food acquisition and food utilization in the particular cultures studied (Mueller, 1990; Joshi and Mueller, 1996).

###### End stage related traits

Surviving individuals in the shallow-type simulated culture had lower minimum critical mass than the dying ones (fig. S9.a.i), whereas no clear pattern emerged for the deep-type culture (fig. S9.a.ii). Till date, no crowding-adapted population from any selection experiment has been shown to evolve reduced minimum critical mass to pupation, but recent studies have indicated that some crowding adapted populations, particularly those reared in cultures similar to the shallow type, may have evolved smaller mass at uncrowded conditions (Sarangi, 2018; Venkitachalam et al., 2023a) – it is yet to be established whether these differences have a link to reduced minimum critical mass.

Both shallow and deep type simulated cultures showed slight differences between surviving and dying individuals in terms of their waste tolerance (fig. S10), with perhaps a greater difference in deep type culture (fig. S10.a). However, the more prominent results were visible in the time dependence of waste tolerance. While early survivors did not have any pattern in terms of their waste tolerance (fig. S10.c), there was a clear linear pattern for the same in dying individuals at later time stages. The early dying individuals had lower waste tolerance, with late dying ones having greater waste tolerance (fig. S10.c). Among these deaths, a few survivors were also present, usually possessing slightly higher waste tolerance than dying larvae at that time step. In the simulations, the larval deaths were exclusively due to high waste concentration, and fig. S10c shows the time at which the waste got intolerable for many small larvae. Of note is also the observation that the larval death occurred earlier and in a smaller time window in the shallow type culture than in the deep type culture (fig. S10c). Experimentally, waste tolerance evolved primarily in populations adapted to cultures resembling the deep type of culture, and particularly in the late eclosing individuals of those populations (Borash et al., 1998; but see Nagarajan, 2009 and Sarangi, 2018 for some contrasting points).

###### Traits from the expanded model – crowding and digging

The deep type simulated cultures had stark differences among survivors and dying individuals in terms of their crowding threshold (fig. S10.5.aii, bii), and slight differences in terms of peak crowding bite rate (fig. S10.6.aii, bii). The survivors appeared to detect crowding at lower thresholds and fed at higher bite rates under crowding compared to those individuals that perished. Smaller differences existed in the shallow type culture in terms of crowding threshold (fig. S10.5.ai, bi), and almost none in terms of peak crowding bite rate (fig. S10.6.ai, bi). This highlights the importance of amelioration of the detrimental effects of crowding in the deep type culture, which faces much higher feeding band densities and thus greater crowding for limited space (fig. 4.c.ii). Surprisingly, in selection experiments, the evolution of increased feeding rate visible as a plastic response to crowding occurred most drastically in populations reared in cultures similar to the shallow type (Sarangi, 2018). We speculate that this could be due to some limit to the plastic increase in feeding rate, which may be capped at lower effective densities, and further crowding would reduce this plastic increase. However, further experimentation on feeding rates at different effective densities is critical to addressing this conundrum.

There was not much difference observable among surviving and dying individuals in terms of their digging length multiplier in the simulated shallow type culture (fig. S10.11.ai, bi). Survivors in the deep culture appeared to have slightly higher respective digging multipliers (fig. S10.11.aii, bii). This is plausible, as digging is likely to be more important in the deep food columns. This difference was probably not as drastic as was seen for other traits, perhaps because larval growth in simulations was a primary determinant of digging length – as long as a larva grew enough to avoid the upper waste-saturated layers, the digging length multiplier would likely only confer slight advantages on average.

##### 4.4.2.3. Exploring population dynamics

We may additionally also carry out various population dynamics experiments using our simulation framework. Unlike in the abovementioned multi-generation extension, each population culture would have a given food volume in a given cylinder, but there would be no control over egg numbers laid each generation. For a given population, the number of eggs laid each generation would instead depend on the number of adults as well as their fecundity (for some such experiments, see Mueller et al., 2000; Dey et al., 2012; Pandey and Joshi, 2022a,b). Recently, it has been observed that populations reared under different combinations of food volumes, total densities and effective densities can evolve different degrees of constancy and persistence stability in their population dynamics (Pandey and Joshi, 2022a,b). Simulations would allow the investigation of dynamics induced in several different types of cultures, which may not be possible empirically due to logistical limitations.

A recent simulation study achieved significant congruence with an experimental population dynamics paradigm (Tung et al., 2019). Survivorship and mass distributions at low and high food levels in the population dynamics experiment were successfully predicted by Tung et al. (2019), based on earlier studies following up on Bakker’s findings (Bakker, 1961; Nunney, 1983; Mueller, 1988b). As their population dynamics experiment was run on set generation times, development time changes due to various egg numbers in limiting food may not be as important in predicting the larval stage outcomes every generation.

Future population dynamics experiments may be carried out to explore if predictions by Tung et al. (2019) can hold in different paradigms exploring various vial surface areas and food column lengths. Integration of some findings of our current study may be important with the earlier model in order to carry out further predictions in more nuanced setups.

In both multi-generation extensions as well as population dynamics simulations from our current study, it may be important to consider a potentially important caveat – even slight deviations from empirical reality in a causal model such as ours could snowball across multiple generations, leading to large deviations when making ultimate predictions. Despite this caveat, however, both the avenues of future studies could respectively direct important experiments for long term selection, or population dynamics, thus leading to a greater understanding of the ecology of competition and the evolution of competitive ability – ultimately rendering the theory of density-dependent selection both more robust and more nuanced.

## Acknowledgments

We thank Profs. Mauro Santos, (UAB, Barcelona) and Sutirth Dey (IISER, Pune) for very helpful discussion regarding the simulation framework. S. Venkitachalam was supported by a doctoral fellowship from the Jawaharlal Nehru Centre for Advanced Scientific Research. This work was supported by a J. C. Bose National Fellowship from the Science and Engineering Research Board, Government of India, to A. Joshi,

## Appendix; Supplementary Material

**Figure S1.**
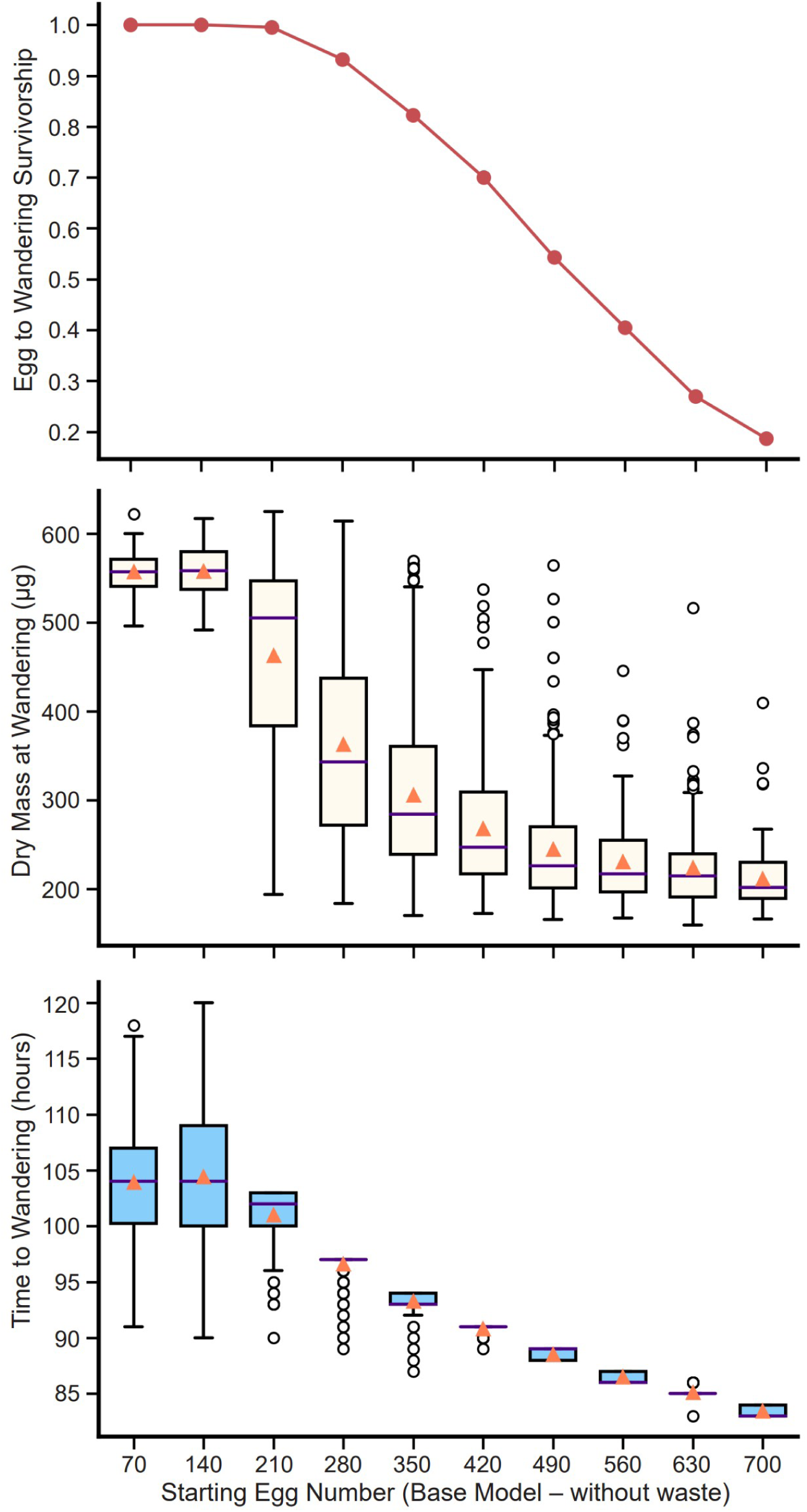
Base model, with larval crowding – waste absent: A single replicate is represented at each point. The X-axis denotes egg numbers. A) Egg to wandering survivorship across increasing egg numbers. B) The distribution of dry mass at wandering (μg) across increasing starting egg numbers. C) The distribution of time to wandering (hours) across increasing starting egg numbers. For the box plots in B) and C), the line within the box symbolises the median, the orange triangle symbolises the mean.

**Figure S2.**
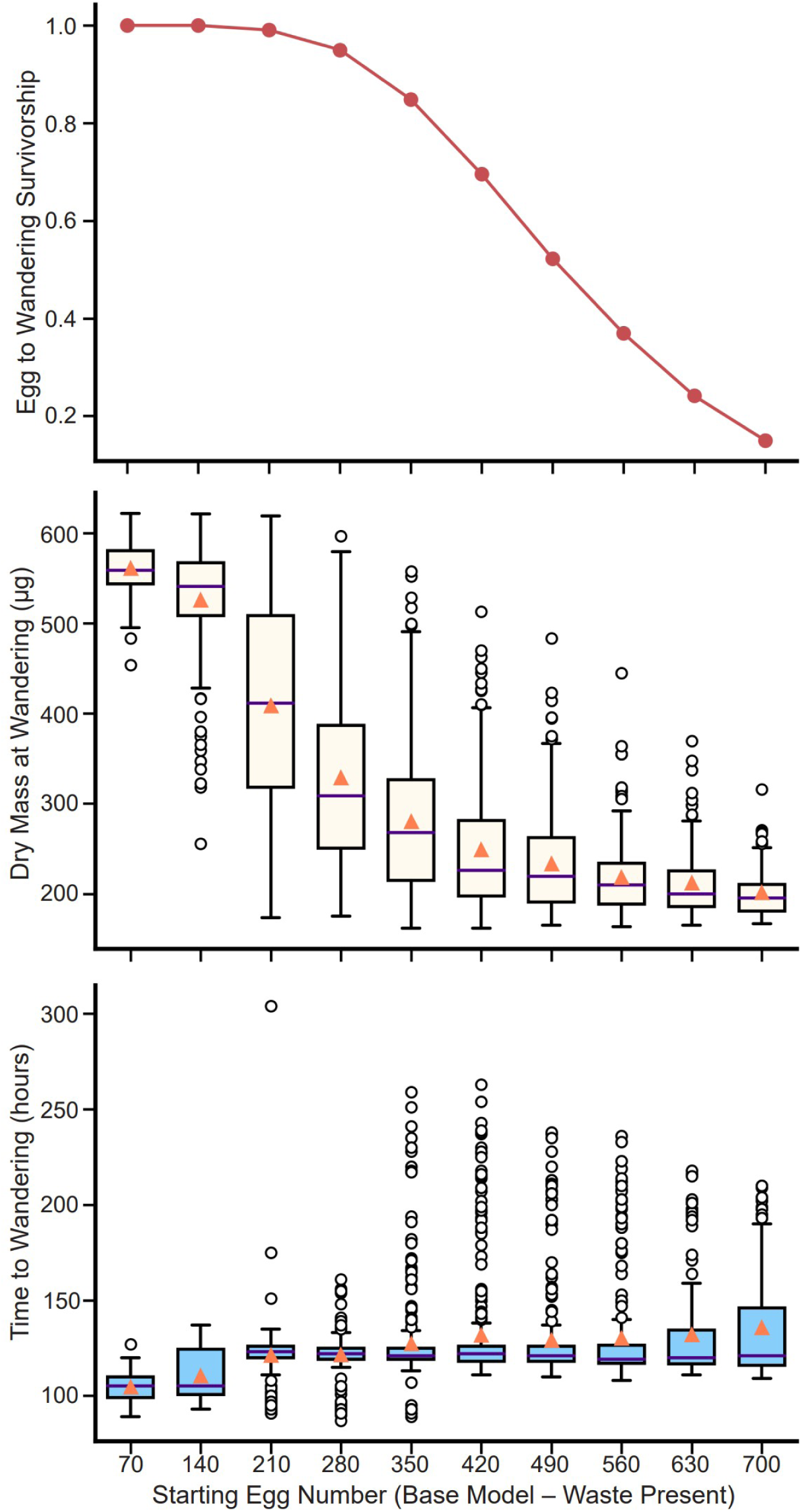
Base model, with larval crowding – waste present. A single replicate is represented at each point. The X-axis denotes egg numbers. A) Egg to wandering survivorship across increasing egg numbers. B) The distribution of dry mass at wandering (μg) across increasing starting egg numbers. C) The distribution of time to wandering (hours) across increasing starting egg numbers. For the box plots in B) and C), the line within the box symbolises the median, the orange triangle symbolises the mean.

**Figure S3.**
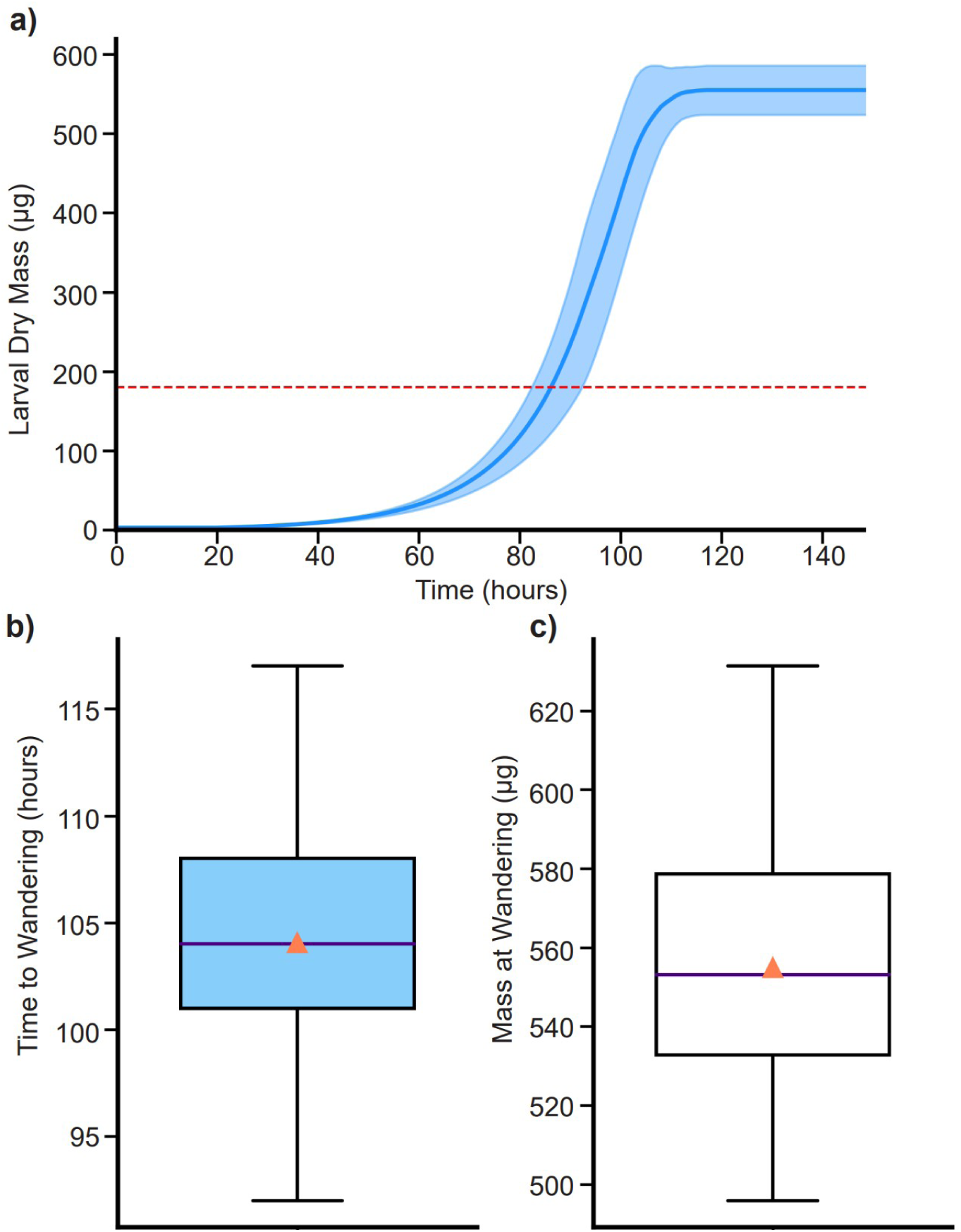
Base model – waste present: Uncrowded (70 eggs in 6 mL food; 400 mm^2^ surface area, 15 mm height). a) Growth profile for 70 individuals. The solid blue line denotes the mean larval dry mass. The shaded region denotes standard deviation in larval dry mass. The dashed red line denotes the mean minimum critical mass to pupation (180 μg). Once the larvae commit to wandering, their mass remains unchanged. b) Time to wandering distribution. The line within the box denotes the median. The orange triangle denotes the mean. c) Mass at wandering distribution. The line within the box denotes the median. The orange triangle denotes the mean.

**Figure S4.**
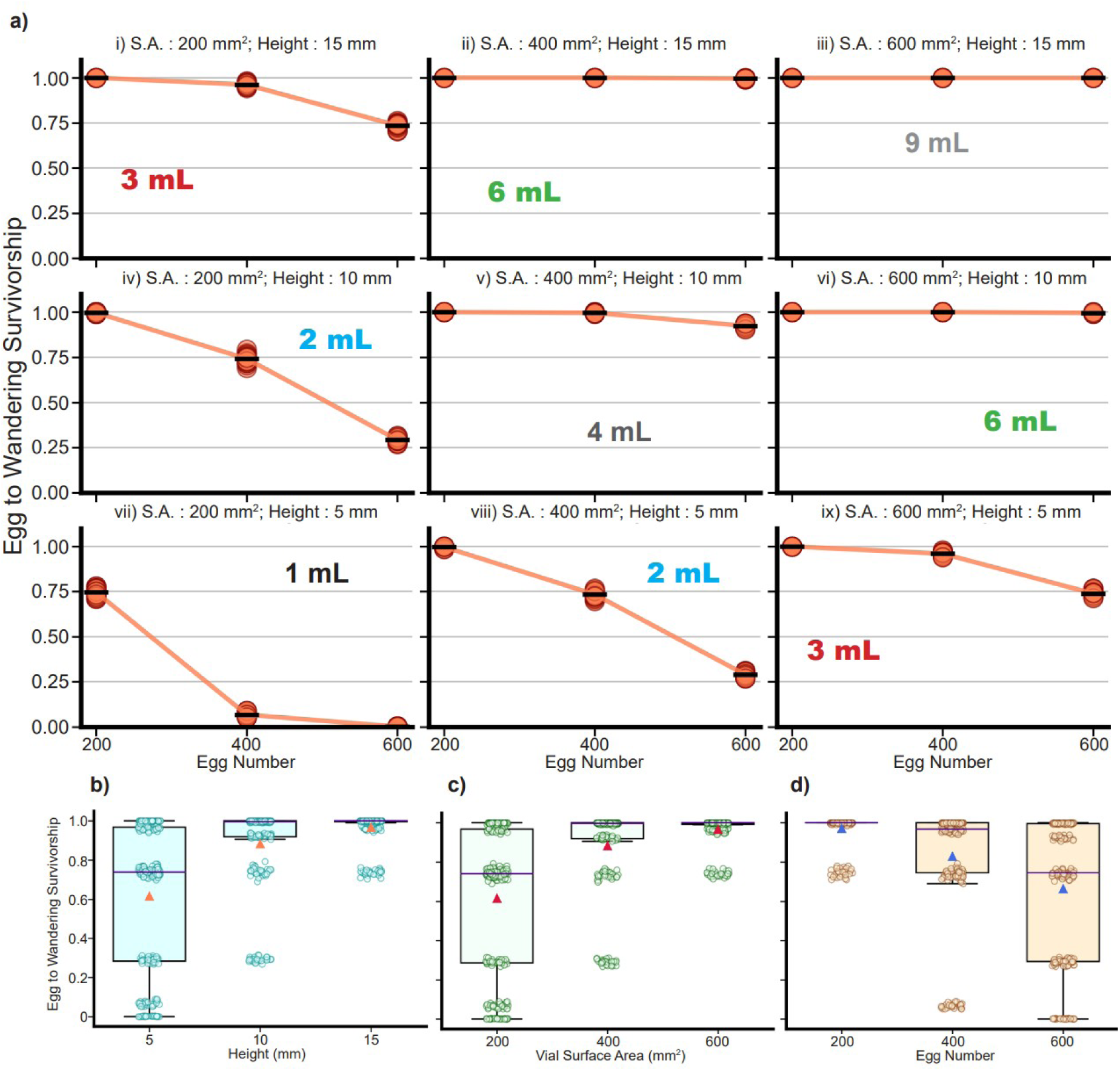
Base model – waste present: Egg to wandering survivorship in a large factorial simulated experiment (see Methods). S.A.: Cross-section surface area of the food column; Height: starting height of the food column. a) Survivorship of 30 replicates for each of the 27 simulated treatments. The ‘-’ symbol in black represents the overall mean for 30 replicates for each treatment. The food volume is also highlighted for each surface area and column height combination. b) Survivorship values for the three levels of food column height pooling data across all other factors. The indigo line within the box represents median, the orange triangle represents the mean. c) Survivorship values for the three levels of surface area of the food column, pooling data across all other factors. The indigo line within the box represents median, the red triangle represents the mean. d) Survivorship values for the three levels of starting egg numbers, pooling data across all other factors. The indigo line within the box represents median, the blue triangle represents the mean.

**Figure S5.**
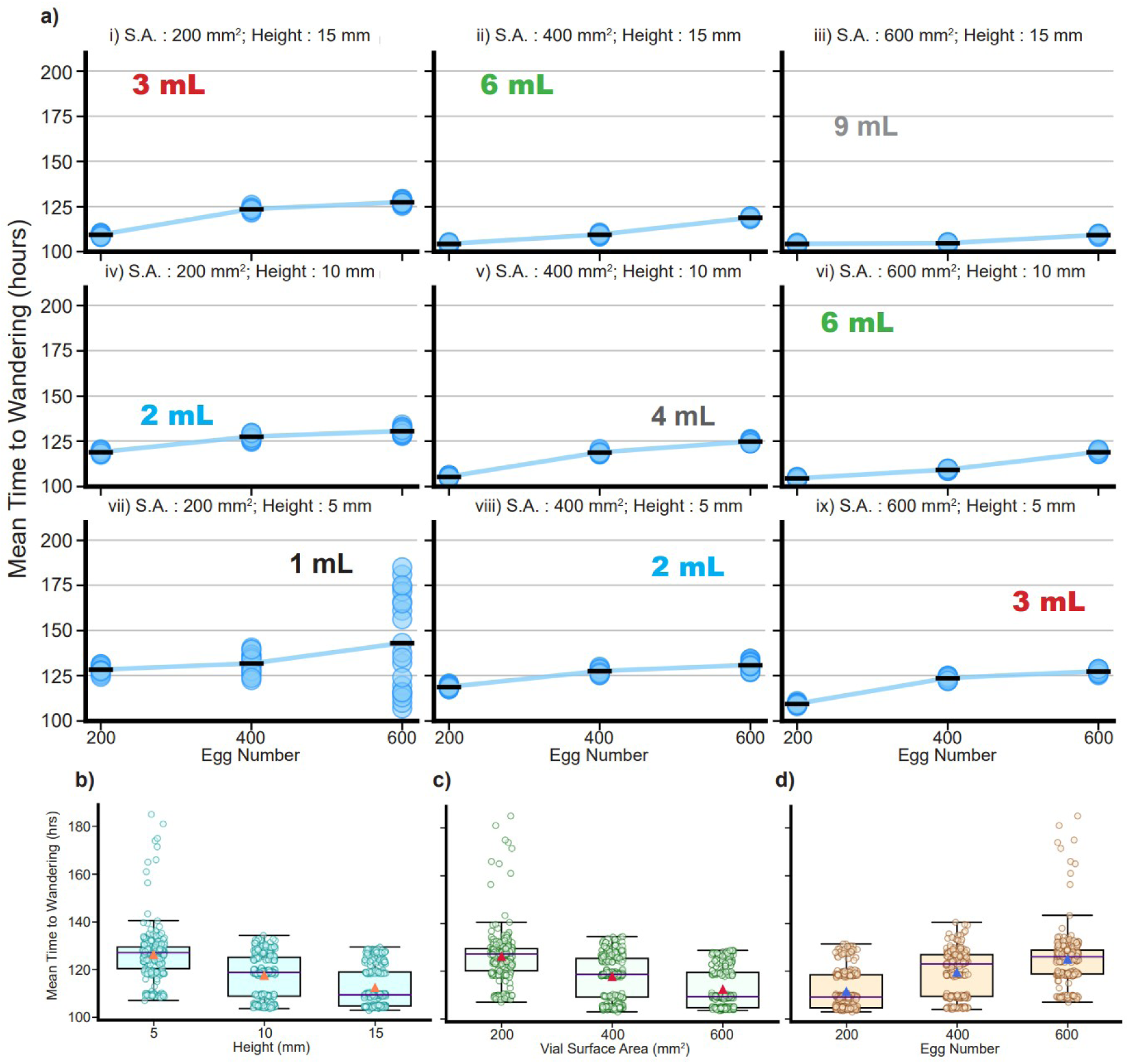
Base model – waste present: Mean time to wandering (hours) in a large factorial simulated experiment (see Methods). S.A.: Cross-section surface area of the food column; Height: starting height of the food column. a) Mean time to wandering of 30 replicates for each of the 27 simulated treatments. The ‘-’ symbol in black represents the overall mean for 30 replicates for each treatment. The food volume is also highlighted for each surface area and column height combination. b) Mean time to wandering values for the three levels of food column height pooling data across all other factors. The indigo line within the box represents median, the orange triangle represents the mean. c) Mean time to wandering values for the three levels of surface area of the food column, pooling data across all other factors. The indigo line within the box represents median, the red triangle represents the mean. d) Mean time to wandering values for the three levels of starting egg numbers, pooling data across all other factors. The indigo line within the box represents median, the blue triangle represents the mean.

**Figure S6.**
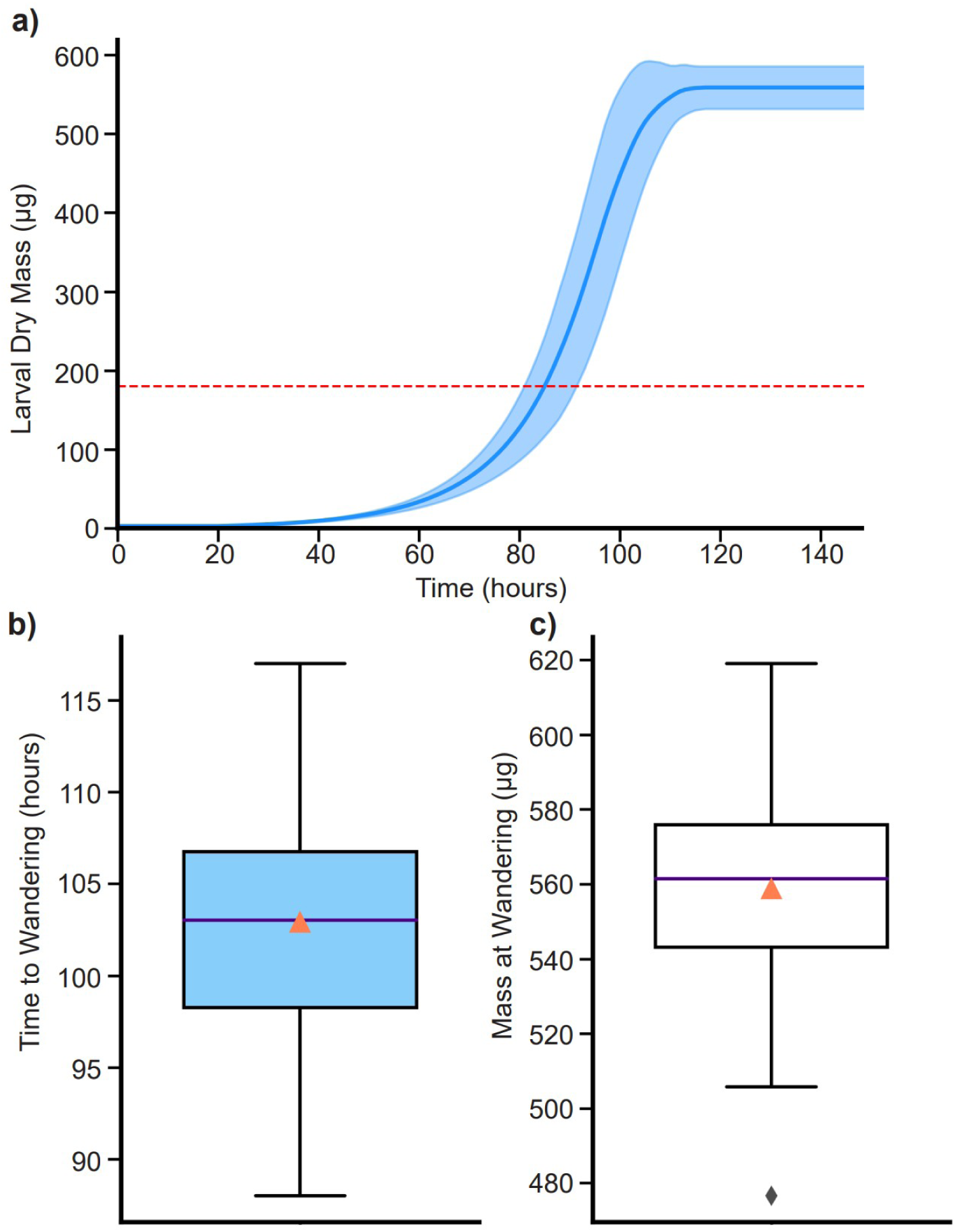
Expanded model: Uncrowded (70 eggs in 6 mL food; 400 mm^2^ surface area, 15 mm height). Growth profile for 70 individuals. The solid blue line denotes the mean larval dry mass. The shaded region denotes standard deviation in larval dry mass. The dashed red line denotes the mean minimum critical mass to pupation (180 μg). Once the larvae commit to wandering, their mass remains unchanged. Time to wandering distribution. The line within the box denotes the median. The orange triangle denotes the mean. Mass at wandering distribution. The line within the box denotes the median. The orange triangle denotes the mean.

**Figure S7.**
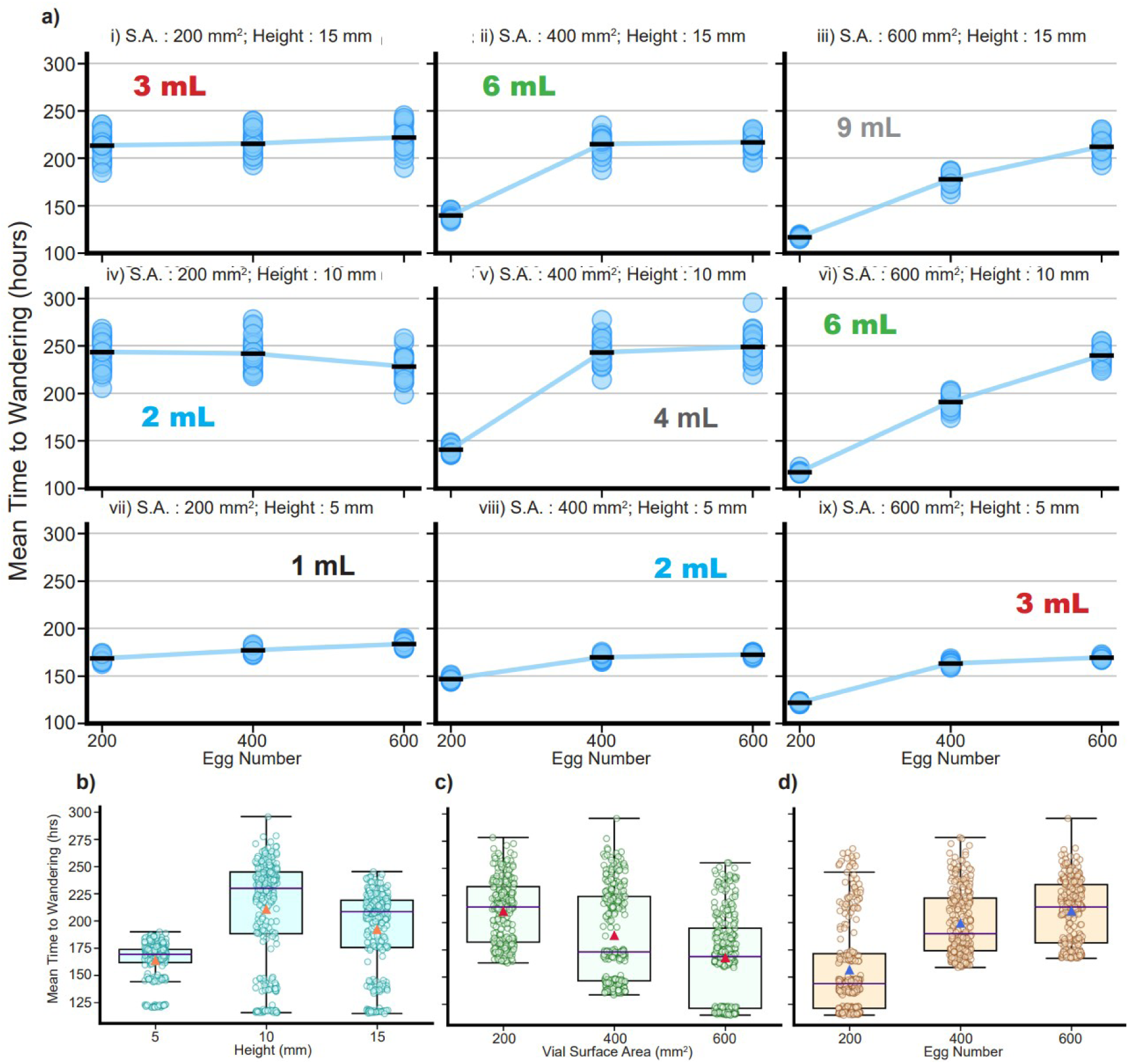
Expanded Model: Mean time to wandering (hours) in a large factorial simulated experiment (see Methods). S.A.: Cross-section surface area of the food column; Height: starting height of the food column. a) Mean time to wandering of 30 replicates for each of the 27 simulated treatments. The ‘-’ symbol in black represents the overall mean for 30 replicates for each treatment. The food volume is also highlighted for each surface area and column height combination. b) Mean time to wandering values for the three levels of food column height pooling data across all other factors. The indigo line within the box represents median, the orange triangle represents the mean. c) Mean time to wandering values for the three levels of surface area of the food column, pooling data across all other factors. The indigo line within the box represents median, the red triangle represents the mean. d) Mean time to wandering values for the three levels of starting egg numbers, pooling data across all other factors. The indigo line within the box represents median, the blue triangle represents the mean.

**Figure S8.**
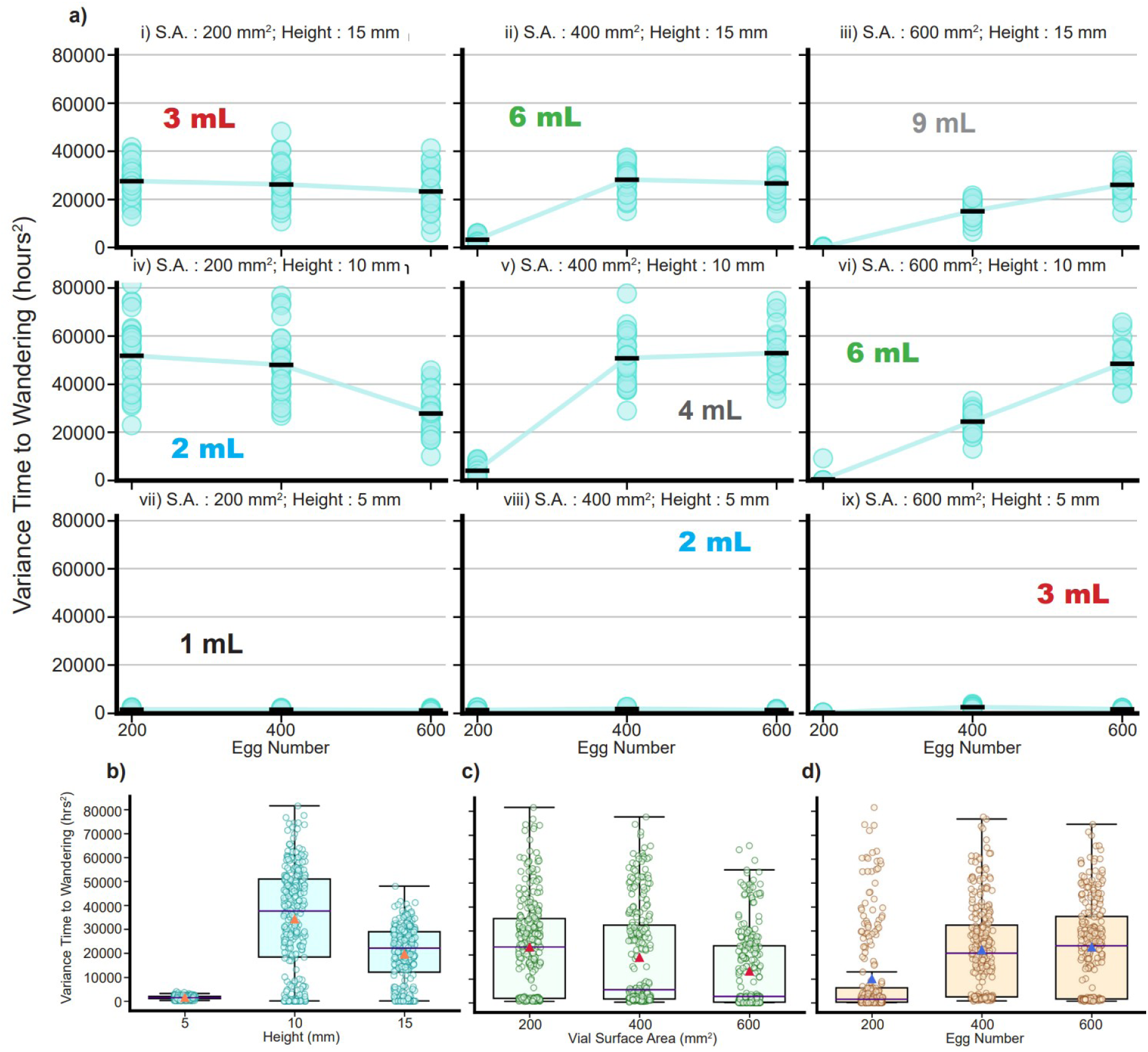
Expanded Model: Variance in time to wandering (hrs) in a large factorial simulated experiment (see Methods). S.A.: Cross-section surface area of the food column; Height: starting height of the food column. a) Variance in time to wandering of 30 replicates for each of the 27 simulated treatments. The ‘-’ symbol in black represents the overall mean for 30 replicates for each treatment. The food volume is also highlighted for each surface area and column height combination. b) Variance in time to wandering values for the three levels of food column height pooling data across all other factors. The indigo line within the box represents median, the orange triangle represents the mean. c) Variance in time to wandering values for the three levels of surface area of the food column, pooling data across all other factors. The indigo line within the box represents median, the red triangle represents the mean. d) Variance in time to wandering values for the three levels of starting egg numbers, pooling data across all other factors. The indigo line within the box represents median, the blue triangle represents the mean.

**Figure S9.**
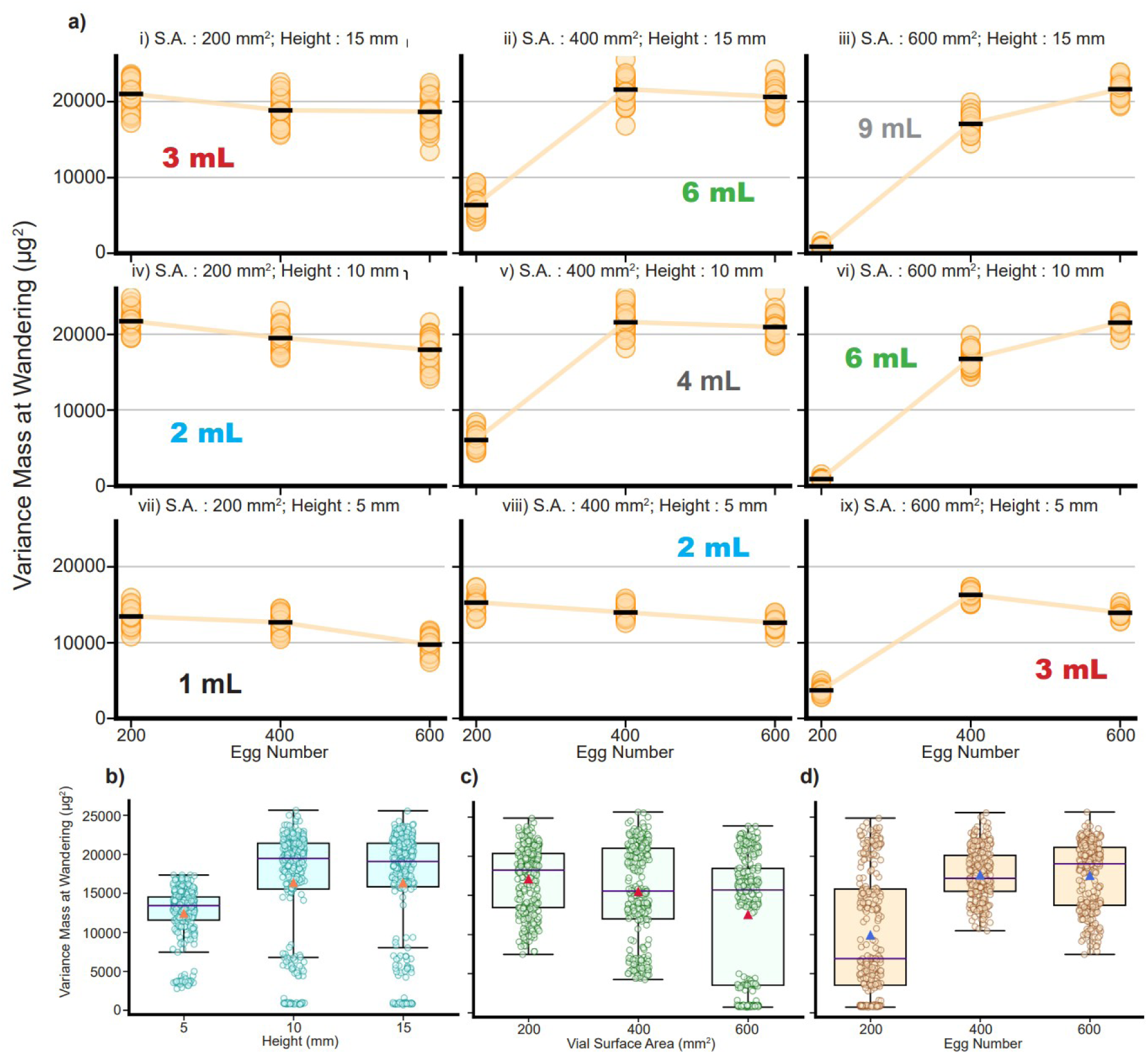
Expanded Model: Variance in mass at wandering in a large factorial simulated experiment (see Methods). S.A.: Cross-section surface area of the food column; Height: starting height of the food column. a. Variance in mass at wandering of 30 replicates for each of the 27 simulated treatments. The ‘-’ symbol in black represents the overall mean for 30 replicates for each treatment. The food volume is also highlighted for each surface area and column height combination. b. Variance in mass at wandering values for the three levels of food column height pooling data across all other factors. The indigo line within the box represents median, the orange triangle represents the mean. c. Variance in mass at wandering values for the three levels of surface area of the food column, pooling data across all other factors. The indigo line within the box represents median, the red triangle represents the mean. d. Variance in mass at wandering values for the three levels of starting egg numbers, pooling data across all other factors. The indigo line within the box represents median, the blue triangle represents the mean.

**S10.**
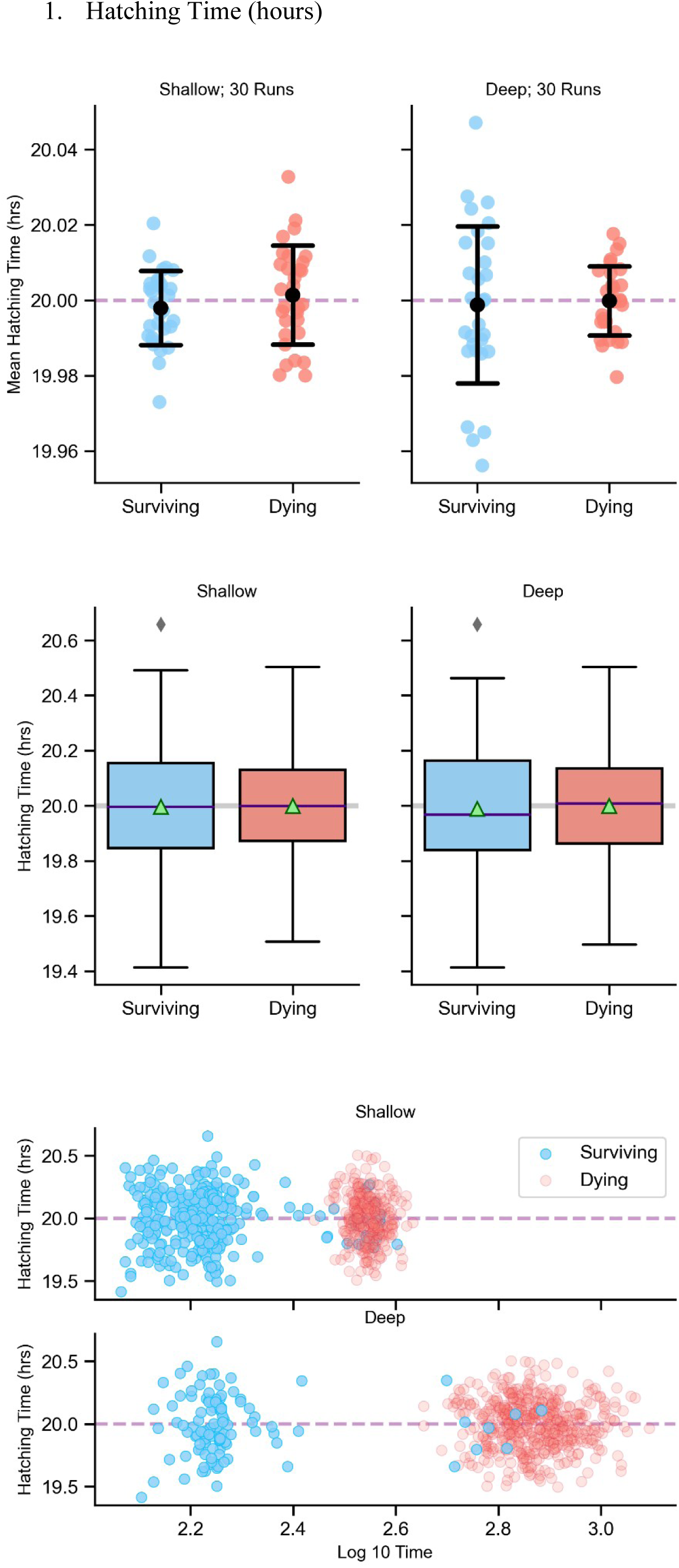
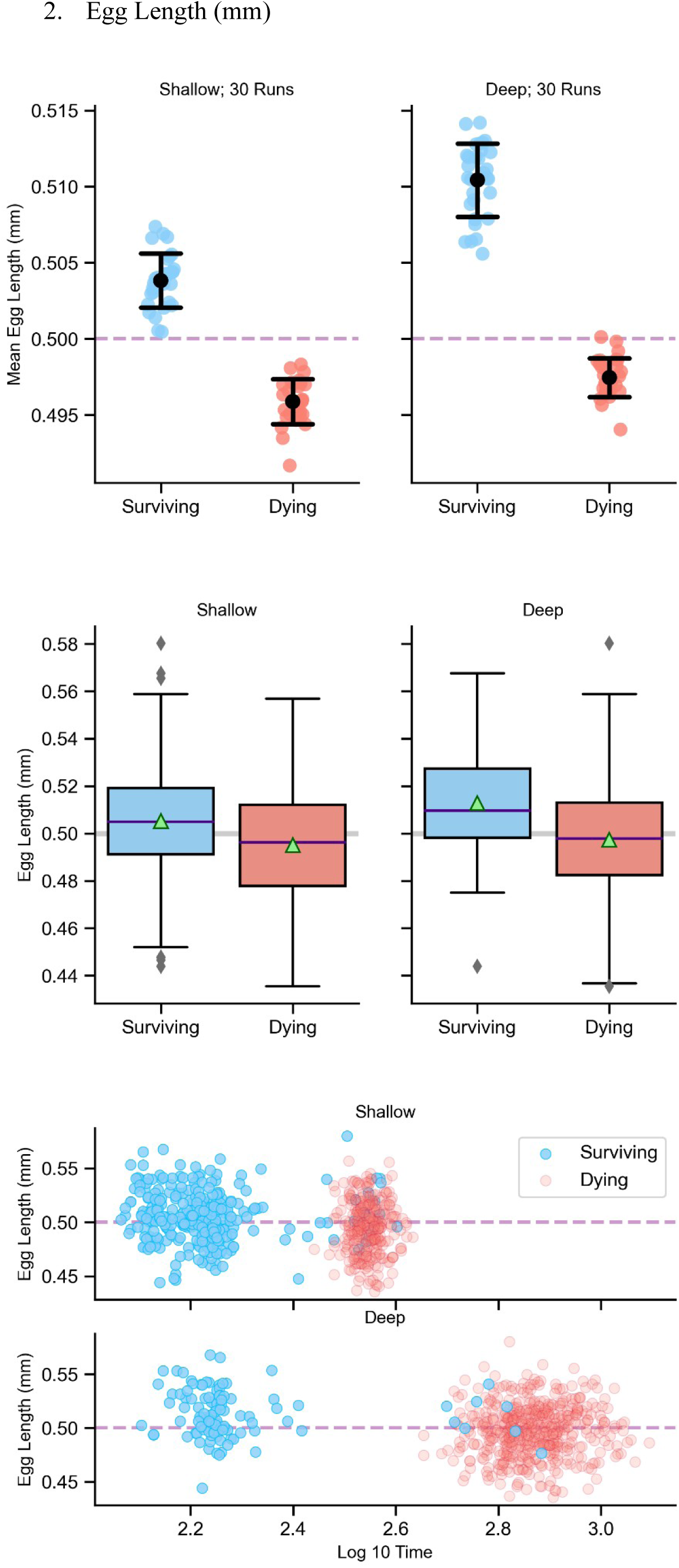
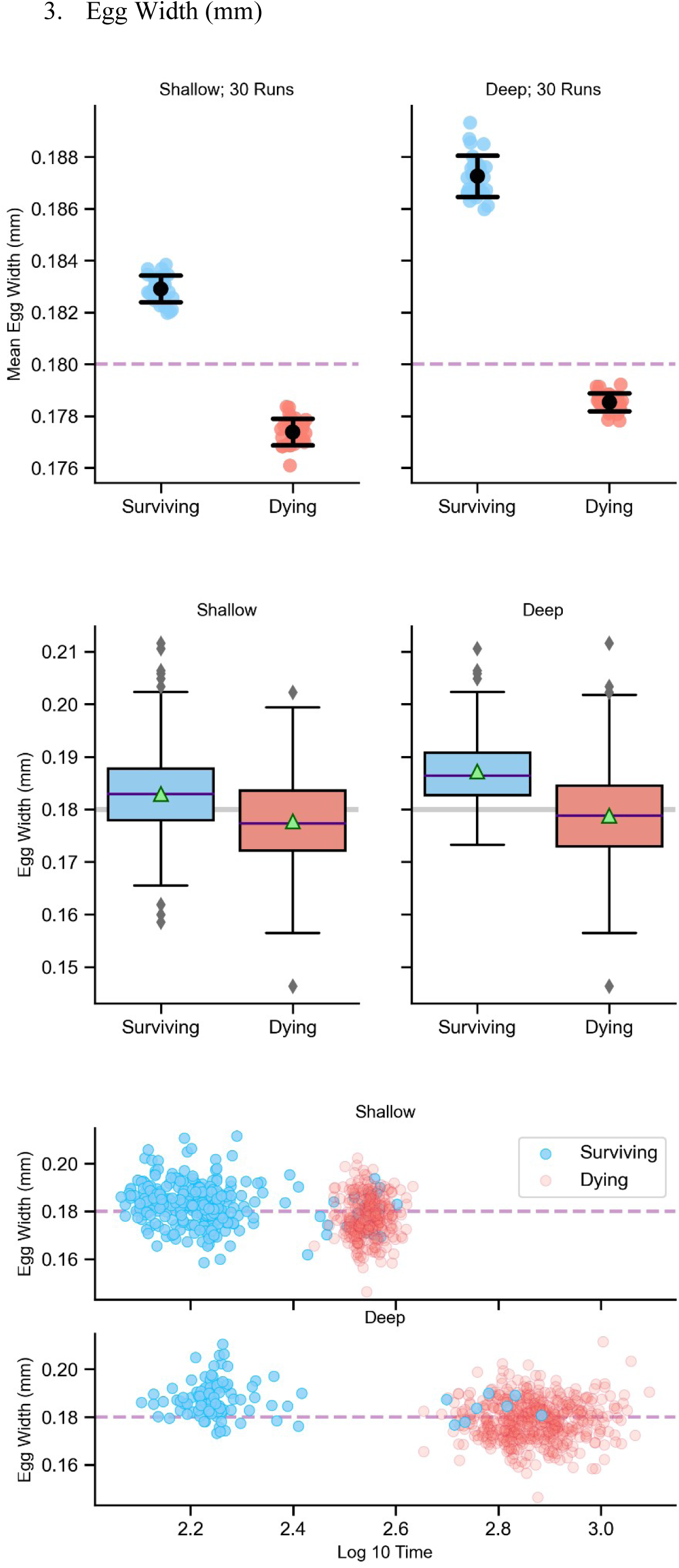
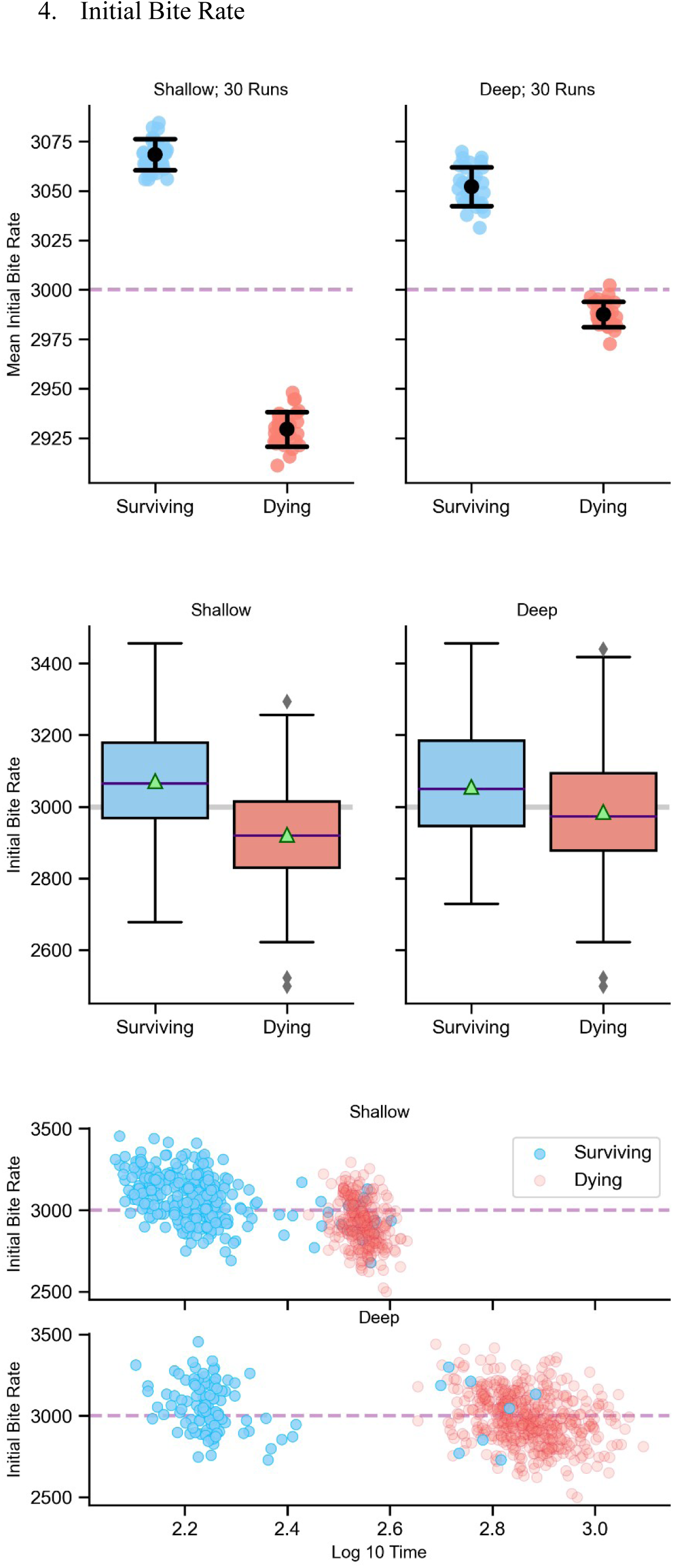
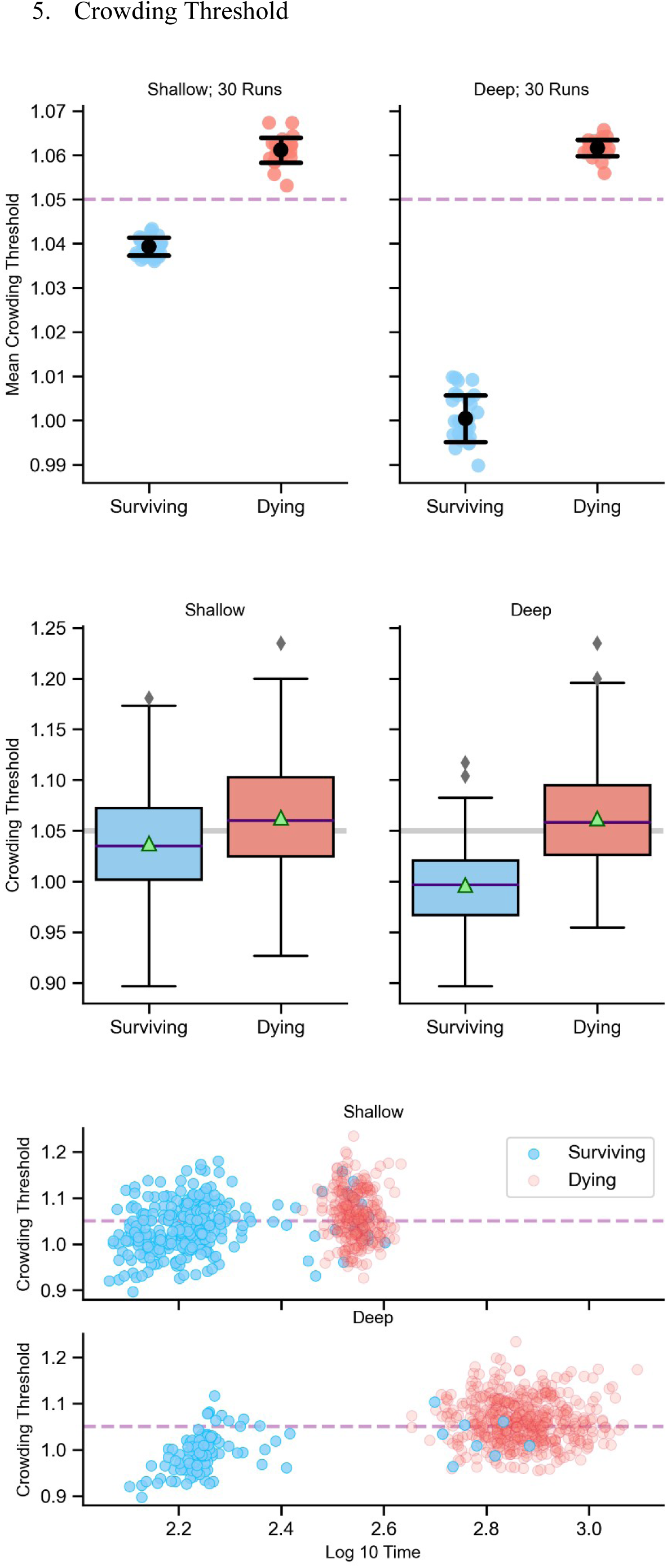
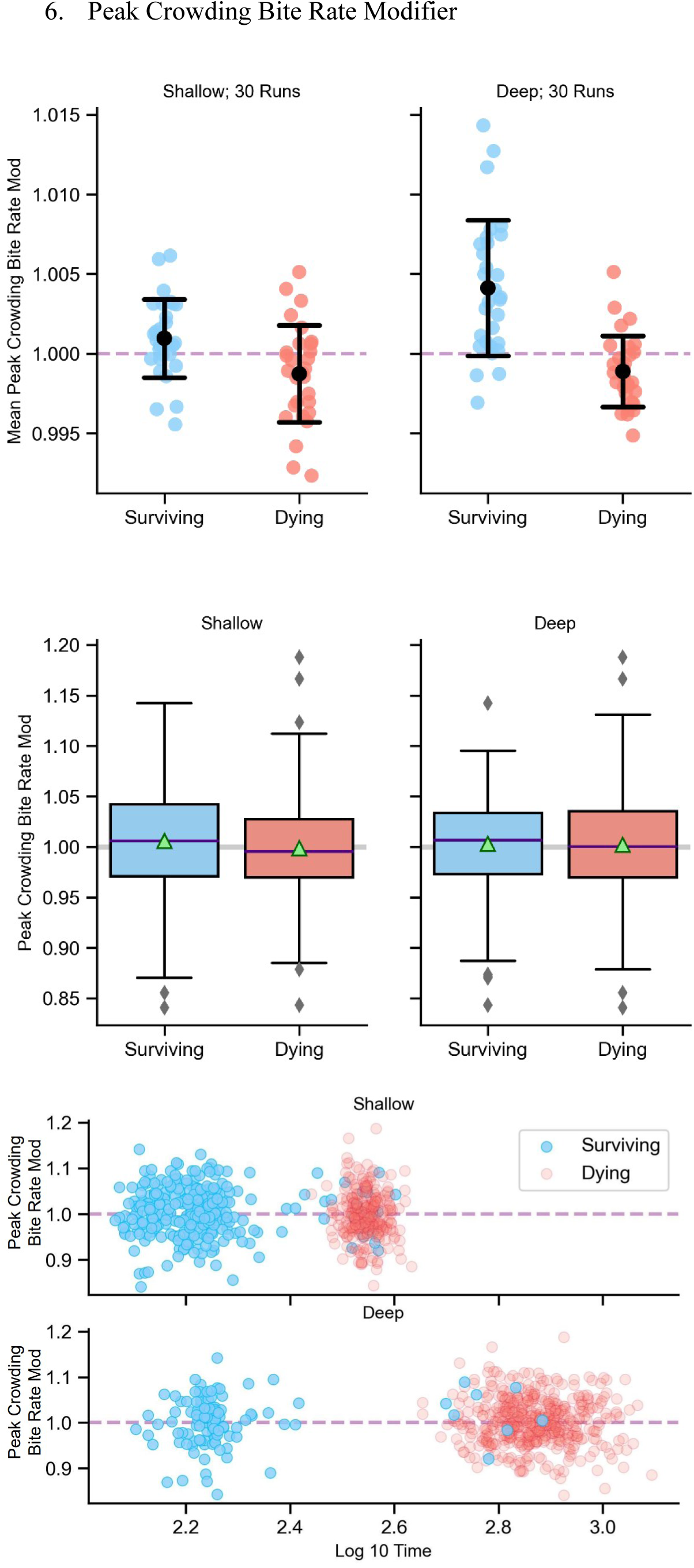
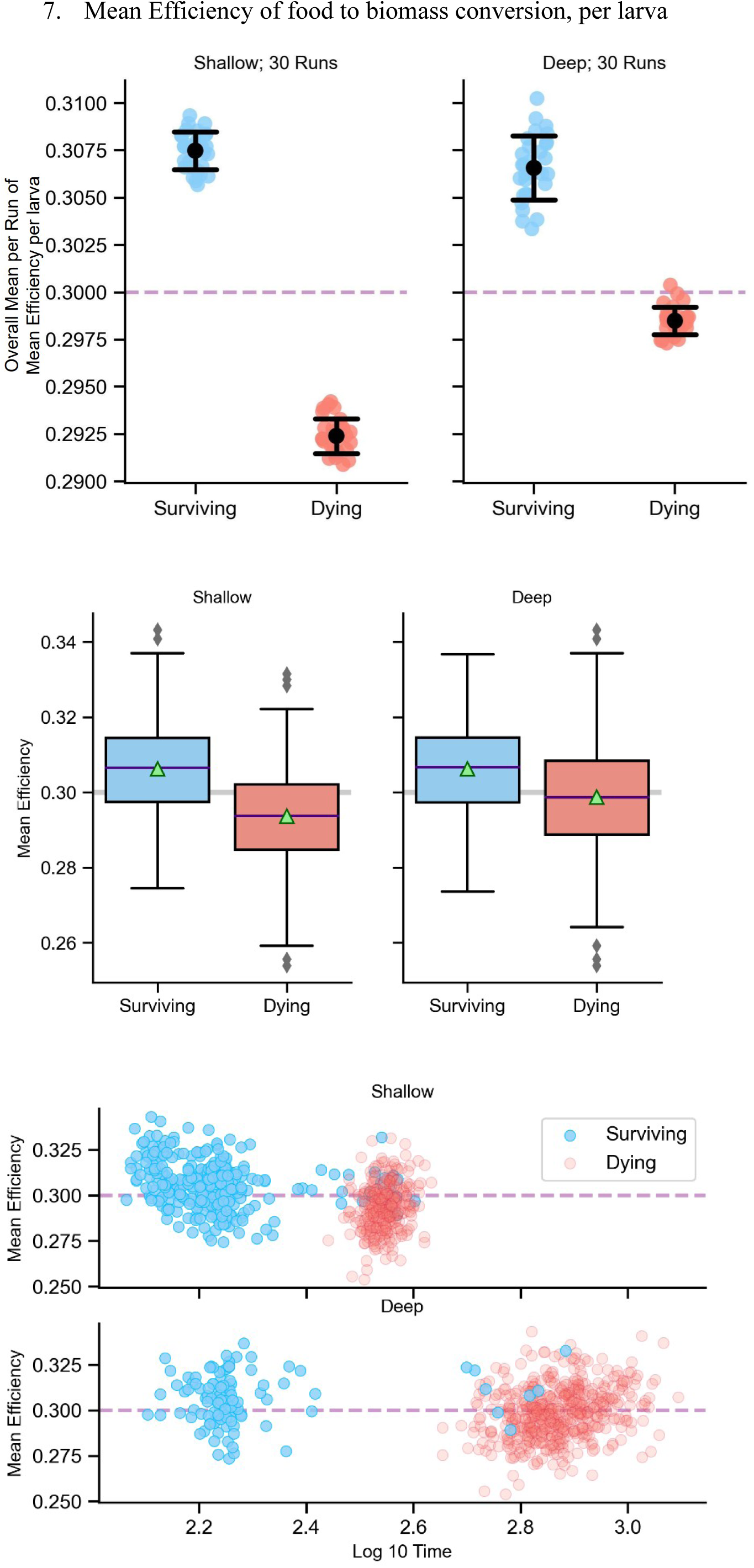
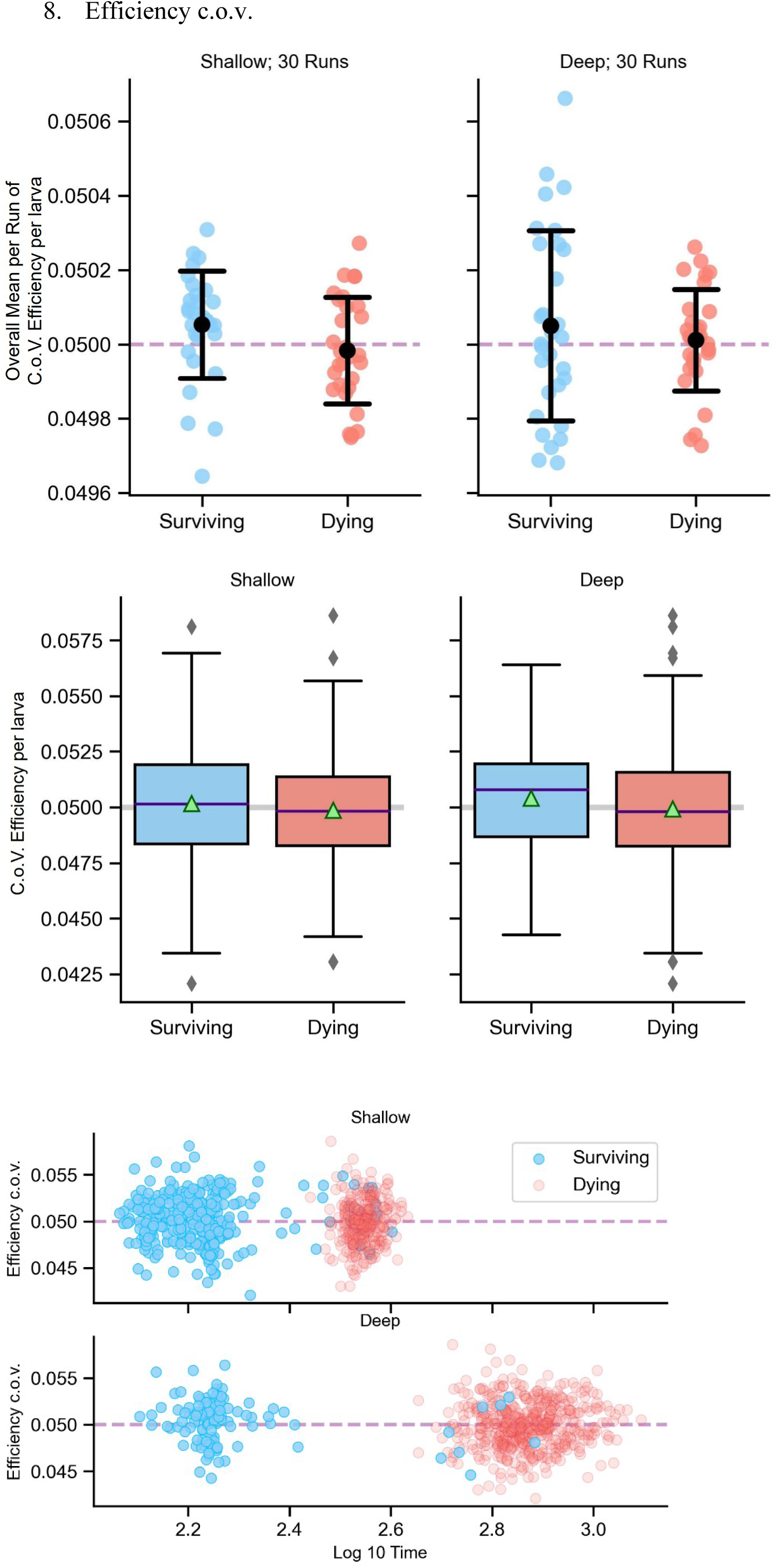
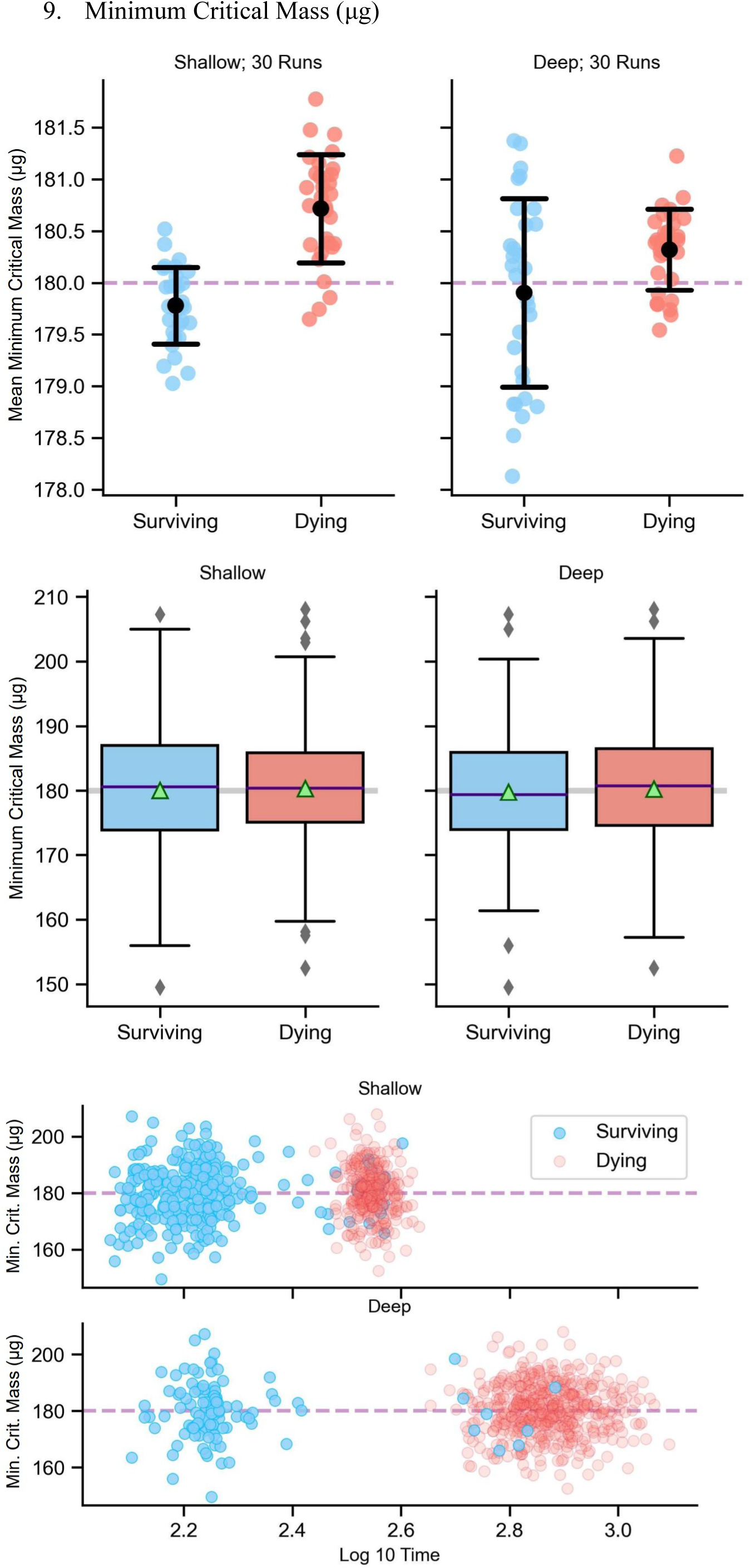
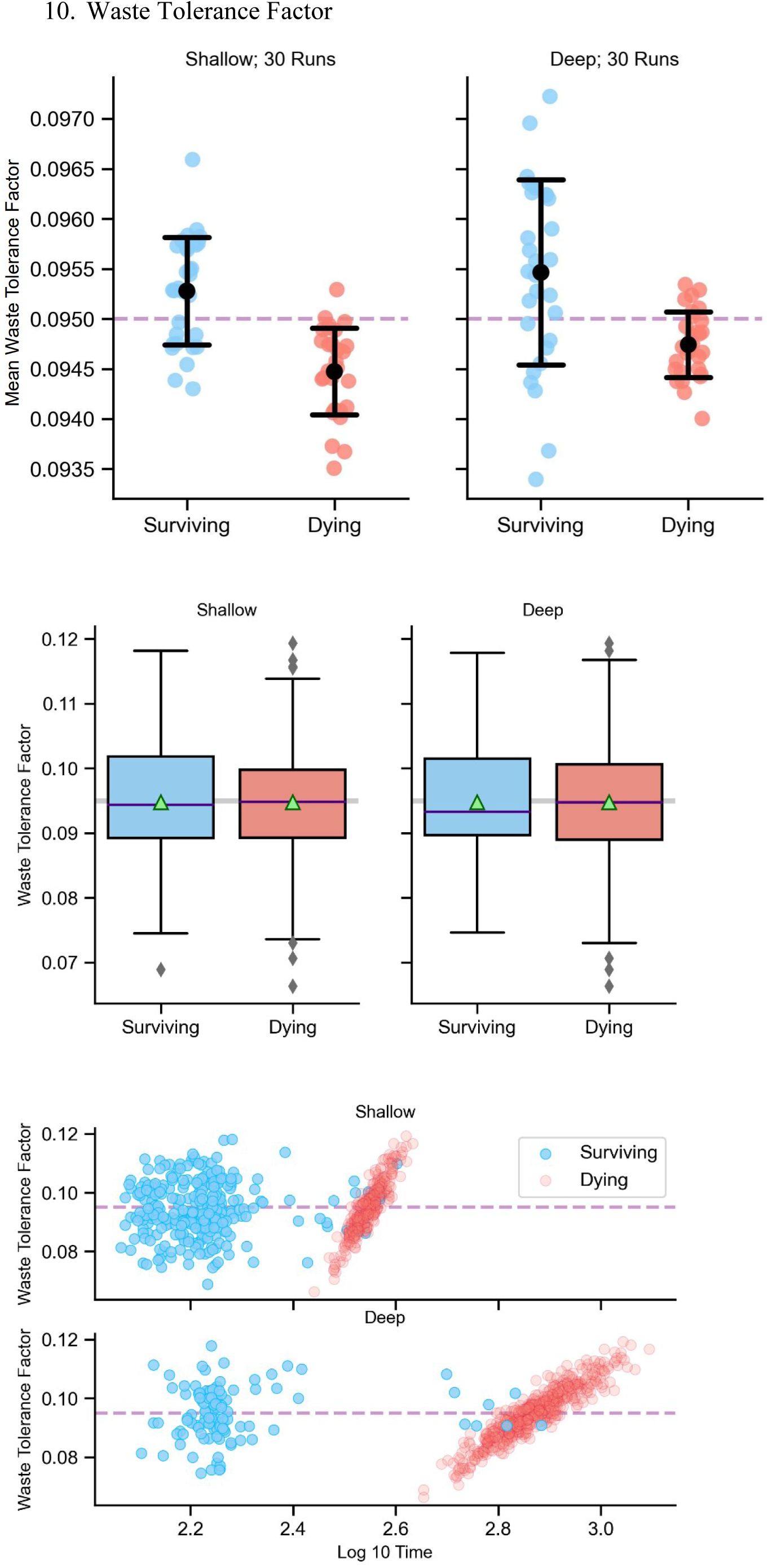
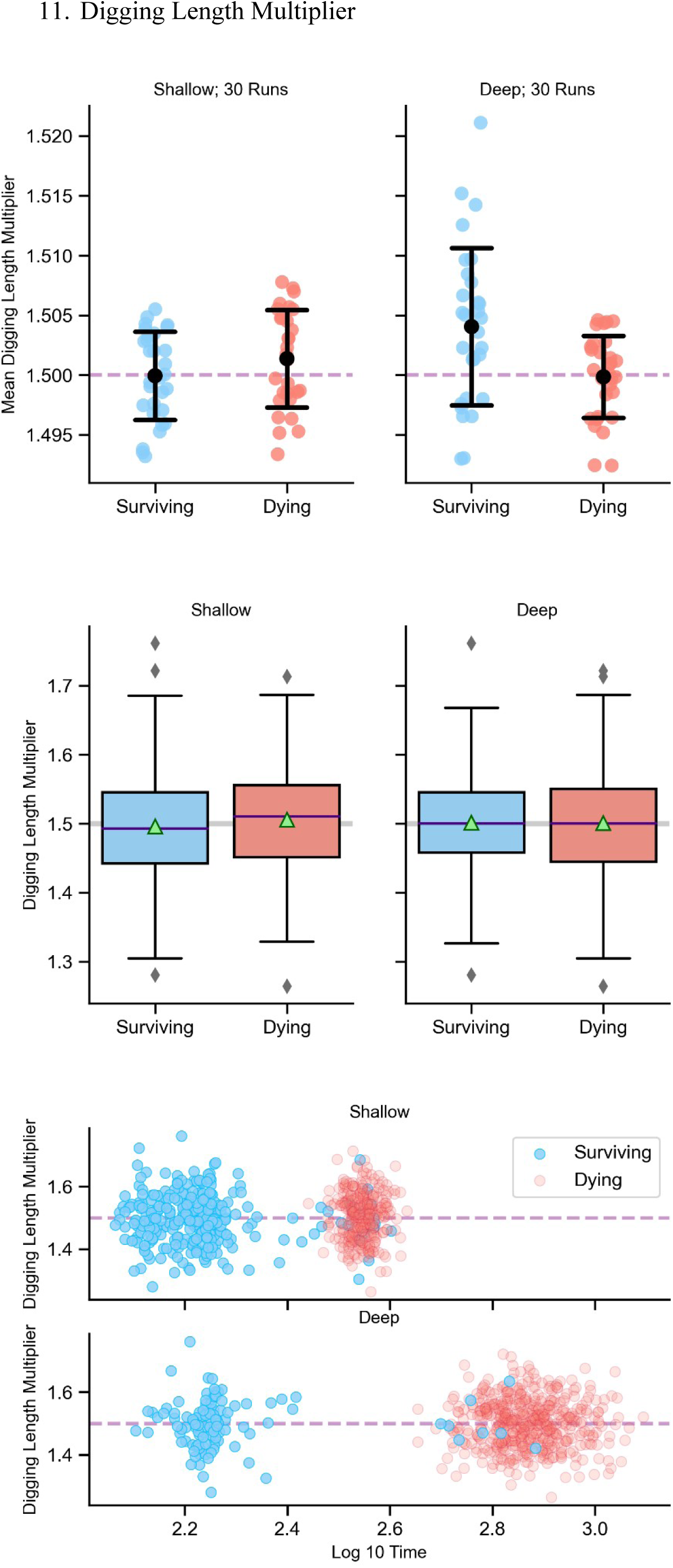
Trait distributions of surviving (blue) and dying (red) larvae: Each numbered figure set represents a trait. The left column of plots represents trait distributions from shallow type cultures – 600 eggs in 3 mL food cast in cylinders of 600 mm2 cross-section surface area and 5 mm column height. The right column of plots represents trait distributions from shallow type cultures – 600 eggs in 3 mL food cast in cylinders of 200 mm^2^ cross-section surface area and 15 mm column height. The first figure in each numbered set represents the mean trait distribution across 30 replicates for surviving and dying individuals, in shallow or deep type cultures. For each replicate, the starting individuals in both types of cultures are the same. Error bars represent standard deviation. The second figure represents the trait distribution of surviving and dying individuals within a single run. The line within the box plots represents the median for each distribution. The green triangle represents the mean for the respective distribution. The third figure represents the trait distribution of individuals across their respective times of wandering (blue) or dying (red). For each trait, the mean value of the distribution from which the traits were randomly drawn at the beginning of the simulation, is also plotted as a purple dashed line (first and third figure per numbered set), or as a grey solid line (second figure per numbered set).

## Notes

### Competing Interest Statement

The authors have declared no competing interest.

